# Modulation of specific interactions within a viral fusion protein predicted from machine learning blocks membrane fusion

**DOI:** 10.1101/2025.09.01.673551

**Authors:** Ryan E. Odstrcil, Albina O. Makio, McKenna A. Hull, Prashanta Dutta, Anthony V. Nicola, Jin Liu

## Abstract

Enveloped viruses must enter host cells to initiate infections through a fusion process, during which the fusion proteins undergo significant and complex structural changes from pre-fusion to post-fusion conformations. Understanding of the fusion protein conformational stability, rapid and accurate identification of the stabilizing interactions are critically important for inhibiting the infections. Here, we leverage molecular dynamics simulations, novel machine learning models and biological experiments to identify the crucial interactions dictating the structural stability of glycoprotein B (gB), a class III fusion protein. We focused on the interactions between the fusion loops and the membrane proximal region in gB, and a new Q181-R747 polar interaction was identified from our machine learning model as critical in stabilizing the gB pre-fusion conformation. Molecular simulations revealed that mutation of Q181 with proline (Q181P) disrupted the fusion loop secondary structure and reduced gB pre-fusion stability. Experiments were designed to evaluate the impact of the Q181P on fusion. Strikingly, the mutation completely abrogated gB membrane fusion activity. The experiments confirmed the importance of Q181-R747 interaction on fusion, which is consistent with the model predictions. The results deepen our fundamental understanding of the molecular mechanisms of gB during viral fusion, which may lead to novel antiviral interventions. The modeling and experimental framework can be generalized to rapidly identify the critical inter-molecular interactions in other important biological processes.

## Introduction

Virus infections remain major threats to human health worldwide. Viruses are obligatory intracellular parasites, and they need to enter the host cells to initiate infection. Enveloped viruses enter the host cells through a fusion process, which is mediated by viral fusion proteins.^1-3^ During the fusion process, the fusion protein undergoes significant and complex conformational changes, which is dictated by complicated molecular interactions between viral fusion protein and other proteins. Glycoprotein B (gB) is a class III (the least well-understood class) herpesvirus fusion protein,^4-6^ and gB is critical for the fusion of many viruses. These viruses include the human pathogens such as herpes simplex virus 1 (HSV-1), HSV-2, varicella zoster virus (VZV), Epstein-Barr virus (EBV), and human cytomegalovirus (HCMV). The implications of these viruses on human health are significant. For example, HSV-1 remains an important human pathogen with a prevalence level of about 67%^7^ and can cause varied outcomes based on the interaction between HSV-1 and the host immune system.^8-11^ HCMV is a leading viral cause of birth defects in developed countries.

In addition to gB, HSV-1 fusion also requires the coordinated interaction of multiple envelope glycoproteins, glycoprotein D (gD), the gH/gL heterodimer.^12^ Upon receiving a signal from gH/gL, gB inserts its fusion loops into the host cell membrane and then refolds to bring the viral and cell membranes together.^13-15^ Endosomal low pH can trigger fusion-associated conformational changes in gB.^16^ gB begins in a meta-stable pre-fusion conformation, and through drastic structural rearrangements during fusion, ends up in a stable post-fusion conformation. The refolding of multiple gB trimers likely creates a pore in the membrane that allows the viral capsid to enter the cell and transport to the nucleus. Clearly, the stability of the gB pre-fusion conformation and the structural changes of gB during the transition to post-fusion conformation are significantly important to the fusion process. Inhibition of “triggers” of conformation changes could block fusion, similar to the case of respiratory syncytial virus (RSV).^17^ Due to the high conservation of gB, investigation of its stability of the pre-fusion conformation could potentially lead to inhibition strategies to all members of the herpesvirus family. Therefore in this work, we aim to elucidate the stabilizing interactions in gB that could be modulated to prevent membrane fusion by altering the relative stability of either the pre-fusion conformation or the post-fusion conformation.

Prior research has indicated important intramolecular gB interactions. For example, the gB cytoplasmic domain (CTD) is thought to negatively regulate fusion^18^ and may act as a clamp that restrains the pre-fusion gB conformation.^19^ The membrane-proximal region (MPR) – which has 40% conserved residues among alphaherpesviruses – can render viruses with negligible infectivity when mutated^20^ and may regulate the mixing of the gB fusion loops with membrane lipids.^21, 22^ The gB ectodomain (containing domains I-V) is thought to interact with the MPR and CTD (**Fig. 1(a)**), and play important roles in regulating the gB pre-fusion to post-fusion and subsequent fusion.^23-25^ Due to the relationship between the gB ectodomain, MPR, and CTD, we theorize that the MPR-fusion loop interaction stabilizes the meta-stable pre-fusion conformation. Further, interruption of this interaction could trigger pre-fusion gB to prematurely undergo conformation change, so that when gB interacts with a cell membrane it is incompetent for fusion.

**Fig. 1:**
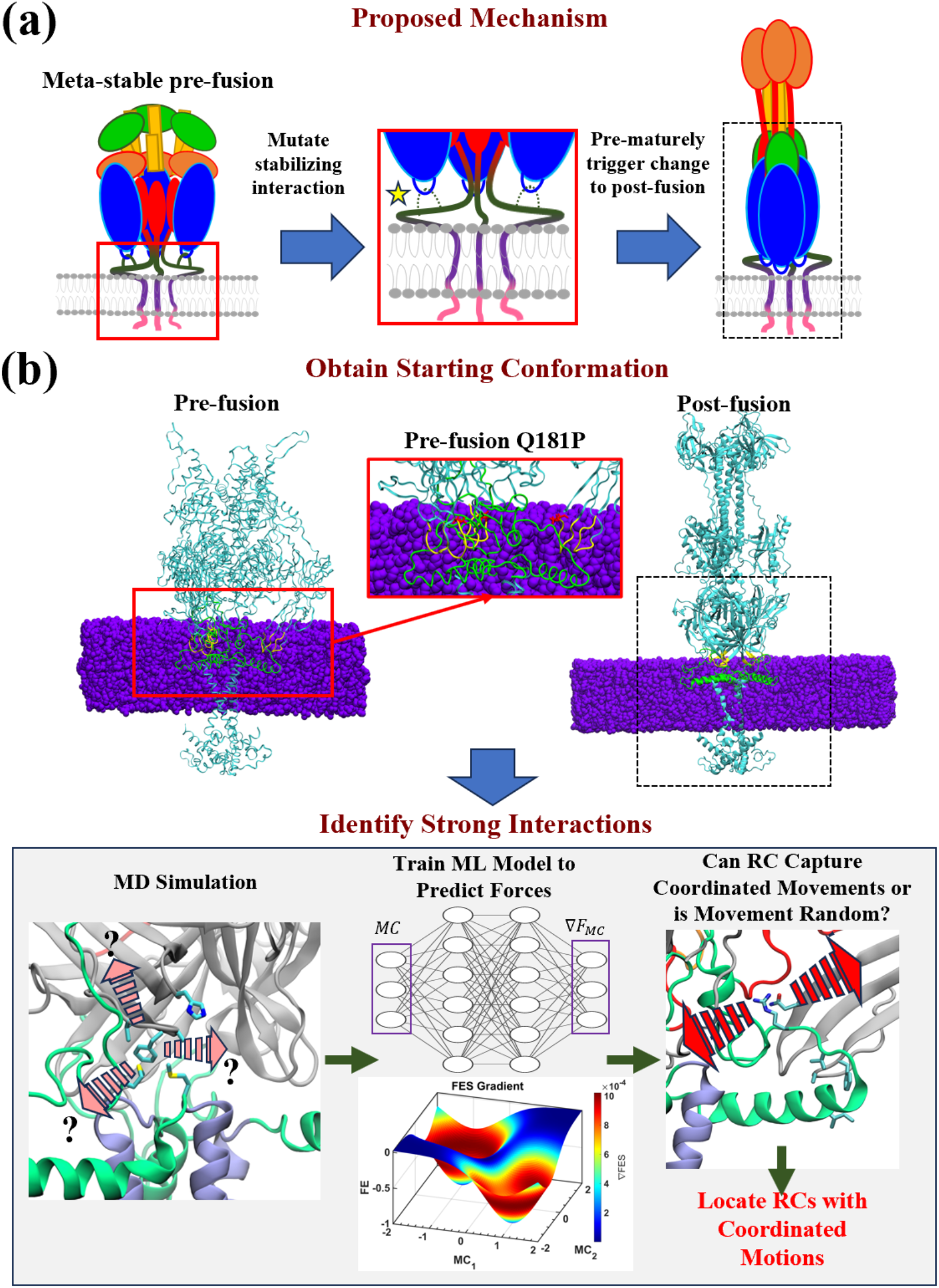
Hypothesis and Computational algorithmic flow. **(a)** The interactions between the fusion loop and MPR are critical in stabilizing meta-stable pre-fusion gB conformation. Destabilization of the critical interactions could trigger pre-fusion gB to prematurely undergo conformation change, so that when gB interacts with a cell membrane, it is incompetent for fusion. The domains I, II, III, IV, and V on gB ectodomain are colored in blue, green, yellow, orange, and red, respectively. The MPR region, transmembrane domain, and cytoplasmic tail domain are colored in dark green, purple, and pink, respectively. **(b)** To prepare the starting pre- and post-fusion conformation, both systems are embedded in a lipid membrane (details can be found in the Method section). To identify the strong critical interactions, an unbiased MD simulation is performed, and user-defined molecular coordinates (MCs) are saved throughout the simulation. A machine learning model is then trained to predict gradients of the free energy surface (forces) and reaction coordinates (RC), which are used to predict an RC for each fusion loop residue in gB. The interaction of each fusion loop residue is evaluated and coordinated interactions are isolated from “random” motions.

In this work, we hypothesize that the interactions between gB fusion loops and MPR are critical for gB conformational changes and consequent virus-cell fusion process. We focus on the identification of the crucial gB fusion loops-MPR inter-residue interactions, which could be potentially modulated to control fusion. To accomplish this, we used a combination of computational and experimental techniques. Molecular dynamics (MD) simulations were first performed to resolve the atomistic-level interactions with high-dimensional trajectory data. Then a machine learning (ML) model was implemented to analyze the data and describe the reactions that occur during the MD simulations. The novel ML algorithm developed by our group, Log-probability estimation via invertible neural networks for enhanced sampling (LINES)^26-28^, were employed to achieve that. The LINES algorithm excels at identifying reaction pathways by learning free energy surface (FES) gradients. Through our modeling and simulations, a new interaction by one of the fusion loop residues was determined to be critical in maintaining the stability of the gB pre-fusion conformation, and was selected for mutational analysis. Mutation of the predicted residue showed dramatic reduction of gB pre-fusion stability from MD simulations. More strikingly, our membrane fusion experiments demonstrated that this mutation in gB completely abrogated membrane fusion activity. The results suggested that it is possible the wild type fusion loop-MPR interaction stabilizes the meta-stable pre-fusion conformation of gB.

## Results and discussion

### Machine Learning and Algorithmic Flow

**Figure 1** illustrates the overall schematic and algorithmic flow of the work, including our hypothesis, preparation of starting configurations for MD simulations, and the steps of identification of strong interactions through our machine learning method. Detailed descriptions of the MD simulations and machine learning models, as well as the definition of the reaction coordinates can be found in the Method section.

### Conformational Movement

To prepare the full-length gB pre-fusion structure, the ecto-domain of the existing pre-fusion structure and the membrane region of post-fusion structure were aligned and superimposed. A truncated gB post-fusion structure from the full-length structure was generated to reduce computational cost. (See Method section and **Figure S1** in Supporting Information for details.) After equilibration, production MD simulations for the pre- and post-fusion conformations are run for 30 ns each. This simulation duration was selected by measuring the autocorrelation times of each molecular coordinate – for which the maximum observed autocorrelation time was 10 ps. Therefore, interactions or residue motions occur on timescales shorter than 10 ps, and 30 ns should be sufficient to sample inter-residue interactions by saving the molecular coordinates every 1 ps. No significant conformation changes are observed over the 30 ns simulations since the starting and ending structures for both pre- and post-fusion conformations are similar, as depicted in **Figure S2** in Supporting Information. **Figure S3** shows the relative movement of both conformations over the 30 ns. It was observed that the post-fusion conformation is exceedingly stable both for the entire truncated structure and for just the domain I-MPR residues. This is not surprising, since the post-fusion conformation is known to be the most energetically stable. In contrast, inspection of the pre-fusion trajectory changes shows large, oscillating motions of a hinge in the domain III segment near G522, but the structure still retains a stable domain I-MPR interaction. The stable domain I-MPR regions indicate that the procedure used to generate the pre-fusion structure with a transmembrane domain was viable. The oscillations of domain III are also expected, since Vollmer *et al*.^*29*^ could only stabilize the pre-fusion ectodomain by introducing the H516P mutation that “locks” a hinge in domain III. It is worthwhile to note that additional simulations were performed without a lipid membrane, but neither of the conformations exhibited a stable pose and quickly destabilized. This paints the picture that the interactions between the fusion loop residues, the MPR region, and the lipid membrane are very stable and also essential for maintaining the pre- and post-fusion conformations.

### Residue Interaction Strengths

To evaluate the inter-residue interaction strength for gB, we implemented our machine learning model^26-28^ to learn the gradients of the free energy surface (FES) with respect to a set of molecular coordinates. A diffusion model^30, 31^ was adopted in our method to estimate the FES gradients due to its flexibility in neural network architecture. Then the reaction coordinates were computed via eigenvalue decomposition of the gradient-based autocorrelation equation. In this work, the distances between fusion loop residues and MPR residues were defined as molecular coordinates. And as illustrated in **Fig. 2(a)**, we calculated a separate reaction coordinate for each fusion loop residue (see Method section for details). The molecular coordinate coefficients between each fusion loop residue and MPR residue are shown in **Figure S4** and **Figure S5** in Supporting Information for the pre-fusion and post-fusion systems, respectively. However, the magnitude of the coefficients does not necessarily indicate whether two residues interact strongly with each other, but rather hints the location of a strong interaction. For example, the MPR residue with the largest coefficient for the fusion loop residue GLN181 is VAL722, but these residues are separated over 1 nm apart from each other. Alternatively, there is a series of moderately large coefficients from ASP744-VAL752. Inspection of the MD trajectories reveals that GLN181-ARG747 is a persistent interaction between the fusion loop and MPR. Since all the neighboring MPR residues also exhibit little side chain movement, the machine learning model does not differentiate between which specific interaction is the strongest and instead “smears” the interaction across all of the MPR residues. Therefore, an interaction strength metric, coordination of correlations (CC), is introduced and calculated based on the weighted L2-norm of the covariance between a reaction coordinate and each of its associated molecular coordinates (details can be found in the Method section).

**Fig. 2:**
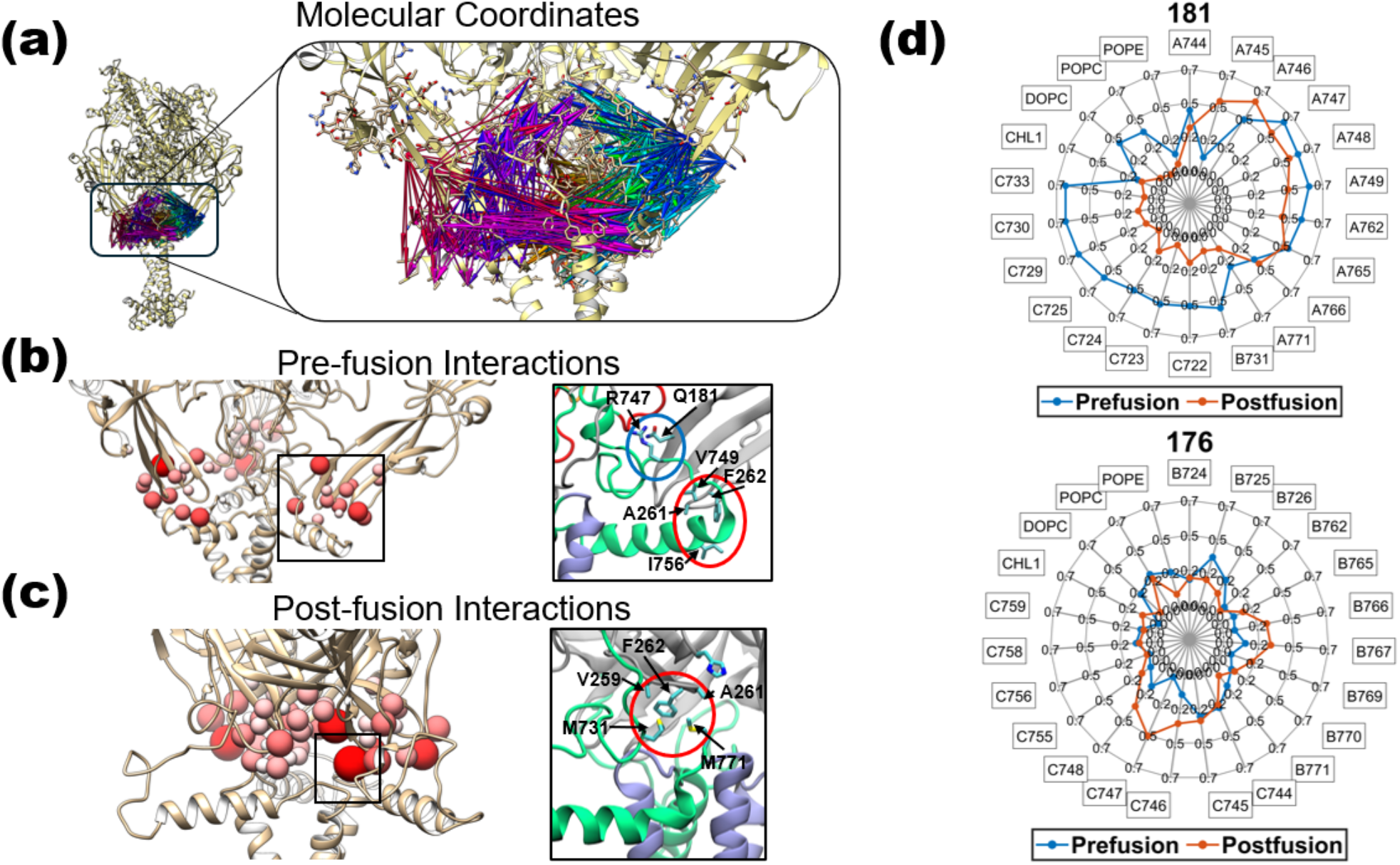
gB computational coordination analysis results. **(a)** The inter-residue distances, colored for each fusion loop residue, are used to predict a separate reaction coordinate for the fusion loop. **(b-c)** The coordinate of correlations (CC) for the pre-fusion and post-fusion interactions are represented by the relative color and size of the spheres around each fusion loop residue, where a redder color and the correspondingly larger sphere radius indicate a greater reaction coordinate coordination. The insets on each conformation represent an example of a strong interaction, where Q181 interacts with R747 in the pre-fusion state and A261/F262 interact with M731/M771 in the post-fusion state. **(d)** The Kendall-Tau correlation coefficient (*τ*) between the reaction coordinate and each MPR residue for Q181 and G176 are compared between the pre-fusion and post-fusion conformations. The Q181-R747 interaction in the pre-fusion state is noted by a relatively long segment of large-value Kendall-Tau correlation, which means many MPR residues are observed to have a coordinated motion with the fusion loop residue. Conversely, G176 does not form any strong MPR interactions.

As shown in **Figure S4** and **Figure S5** in Supporting Information, from the calculated CC values some of the strongest fusion loop-MPR interactions in the prefusion conformations are for Q181, A261, and F262. In the pre-fusion state, Q181 forms polar interactions with R747, shown in **Fig. 2(b)**, while A261 and F262 form a hydrophobic cluster with V749 and I756. A261 and F262 also have some strong interactions in the post-fusion state with M731 and M771, as shown in **Fig. 2(c)**. Interestingly, both residues of A261 and F262 have been previously studied by Hannah *et al*.^32^, and A261 was reported to be essential for gB’s function in cell-cell fusion. Hannah *et al*. also analyzed V259, but the studies did not observe significant impacts on V259R, F262L, or F262D mutations for preventing gB function. However, since the V259 and F262 mutations were not implemented in tandem with each other or with A261, it is possible that the mutations were not able to disrupt the hydrophobic clustering in the pre- or post-fusion conformations, thereby allowing for the hydrophobic cluster to remain and stabilize the conformations. It is worthwhile to note that residue H263 also demonstrated a large value of CC. But visual inspection of the trajectory revealed that H263 interacts with other fusion loop residues, possibly in a stabilizing manner, but does not interact with the MPR. Additionally, the relatively large value of CC in H177 for both pre- and post-fusion conformations is due to the close proximity of H177 to other fusion loop residues V259-F262 that are exceedingly stable.

The importance of looking into the correlations between each fusion loop residue and its neighboring MPR residues in different monomers is illustrated in **Fig. 2(d)**, which shows the Kendall-Tau correlation coefficients (details of calculations can be found in Method section) for Q181 and G176 in both the pre- and post-fusion conformations. For example, Q181 has a series of large correlations in monomer C from residues 729-733. However, these MPR residues are not in direct proximity to one another, which suggests that residues 729-733 may reside in a stable patch of residues and that Q181 may instead have a strong interaction elsewhere in that stable patch, namely, residue 747 of monomer A (see **Figure S1** in Supporting Information for illustration of the three monomers). Conversely, **Fig. 2(d)** also shows the interaction results for G176, a residue which does not show any strong interactions in either the pre- or post-fusion conformations. To get a full picture of what is occurring, one should use the correlations to identify the stable patches and then locate the proximal residues with which an interaction is actually occurring. The marked difference in the magnitudes for Q181 and G176 demonstrates how these CC values can be used to identify important interactions.

### Mutation Simulations of Q181

From the residue interaction strength analysis above, Q181 appears to be a crucial residue that has not been previously studied and warrants further investigation. Since this current work focuses on the interactions between the fusion loop residues and the MPR domain, we will apply further analysis of Q181 and its role in stabilizing the pre-fusion conformation. One mutation that could be potentially introduced in experiments is a proline mutation. We will analyze changes in the Q181 interaction strength for this mutation, referred to hereafter as Q181P.

Following 30 ns of MD simulation, the mutated structure is still in a pre-fusion conformation, as shown in **Figure S6** in Supporting Information. The trajectory is then used to train separate ML models and produce reaction coordinates. The value of CC of the predicted reaction coordinates for capturing reactions is computed over the 30 ns trajectories, and the end state is shown in **Fig. 3(a)**. The mutation is noted to decrease the interaction strength (CC) between the MPR and fusion loops from 0.42 ± 0.08 in the wild type pre-fusion conformation to 0.31 ± 0.14 for the Q181P mutation. Here the error margins come from the standard deviation for the 3 monomers. As shown in **Fig. 3(b)**, the Q181P proline mutation modifies both the side chain and backbone of residue 181, disrupting the secondary structure of the fusion loop and the interactions with R747. This creates a gap between Q181P and other - potentially stabilizing - residues like T745 or R747 that would otherwise have a stabilizing effect when interacting with Q181. A proposed schematic of the mutation impact is shown in **Fig. 3(c)**, where the Q181P mutation disrupts the fusion loop secondary structure and prevents interactions with R747 or other MPR residues that would otherwise stabilize the pre-fusion conformation.

**Fig. 3:**
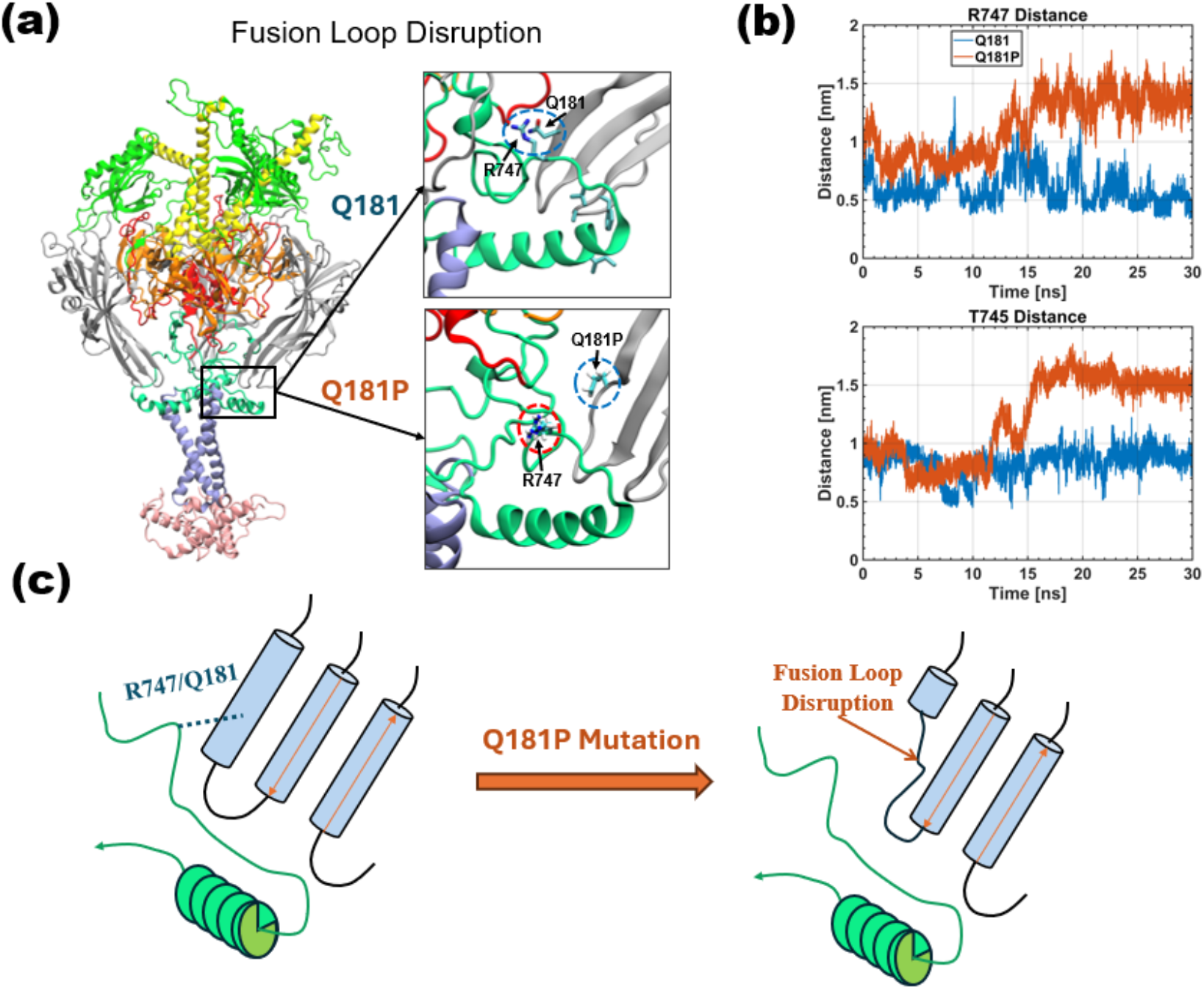
gB Mutation Impacts on Pre-fusion Conformation. **(a)** A depiction of the fusion loop-MPR interaction for the Q181P mutation, with the wild-type results included for reference. Q181P disrupts the fusion loop interaction with R747 and introduces a kink in the fusion loop. **(b)** The Q181P mutation breaks the residue 181 bond with R747 and disrupts the secondary structure of the fusion loop. **(c)** The proposed mechanism is that the wild type gB has a stabilized pre-fusion conformation from the Q181-R747 interaction, and the Q181P mutation disrupts the fusion loop (FL) structure and destabilizes the pre-fusion conformation.

### Q181P mutation abrogates the membrane fusion function of gB

Residue 181 was identified by simulation to be important for stability of the gB pre-fusion conformation. Site-directed mutagenesis is a common molecular biology approach to identify residues important for the structure or function of a wild type protein. We experimentally tested the effects of altering Q181 on the structure and function of gB. The wild type HSV-1 gB gene in a plasmid expression vector (**Figure S7** in Supporting Information) was modified by site-directed mutagenesis, resulting in a gB mutant protein with a substitution of the Q at position 181 with a P (**Figure S8** in Supporting Information). Cells were transfected with the plasmid, and protein expression of gB Q181P was detected by western blot (**Fig. 4(a)**). The HSV-1 gB molecule bearing the Q181P mutation was expressed on the cell surface to near wild type levels as determined by CELISA (**Fig. 4(c)**, bottom panel). This suggests that the Q181P mutation did not grossly affect transport of gB to the plasma membrane and that gB Q181P can be fully evaluated for cell-cell fusion activity, which is dependent on surface expression of gB. We employed a virus-free reporter assay to measure fusion activity of the mutant protein (**Fig. 4(b)**). When cells that have wild type gB gD, gH, and gL on their surface are added to target cells bearing host cell receptors for HSV-1, there is detectable fusion of the two cell populations. Notably, gB Q181P exhibited negligible cell-cell fusion activity compared to wild type gB (**Fig. 4(c)**, top panel). To probe the antigenic structure of the fusion-inactive gB Q181P, we employed a diverse panel of monoclonal antibodies that recognize distinct epitopes all across the gB molecule (**Fig. 4(d)**). As determined by immuno-dotblot, reactivity of antibodies to gB Q181P domains III and VI was similar to wild type gB, while antibodies to domains I, IV, and V exhibited < 1.8-fold difference in reactivity (**Fig. 4(e)**). This suggests that while gB Q181P is functionally inactive for fusion, it retains a conformation that is not grossly altered. These experimental results suggest that gB fusion loop interaction with the membrane proximal domain is critical for maintaining the metastable pre-fusion conformation of gB. A proline residue at position 181 may convert or pre-trigger HSV-1 gB to assume a post-fusion form, rendering it inactive for fusion.

**Fig. 4:**
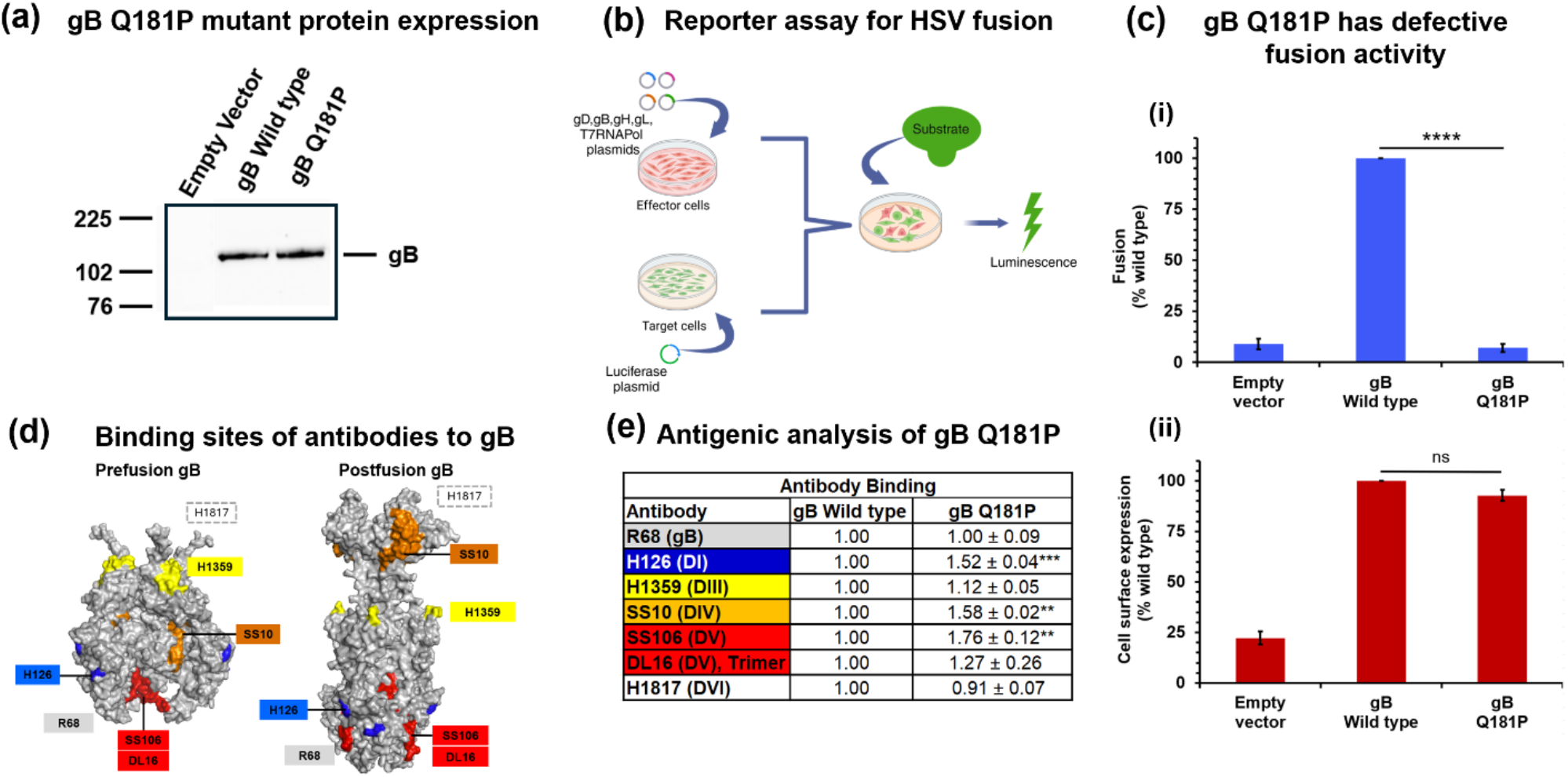
Q181P mutation abrogates the membrane fusion function of gB. **(a)** CHO-K1 cells were transfected with plasmids encoding wild type gB or gB Q181P. Lysates were collected and resolved by SDS-PAGE. Western blot was probed with anti-gB rabbit polyclonal antibody R68. Molecular weight standards (in kilodaltons) are shown to the left. **(b)** A schematic of luciferase reporter assay for cell-cell fusion. CHO-K1 (effector) cells were transfected with plasmids encoding T7 RNA polymerase (pCAGT7), plasmids pPEP98 (wild type gB) or pAM26 (gB Q181P), pPEP99 (wild type gD), pPEP100 (wild type gH), and pPEP101 (wild type gL). CHO-HVEM (target) cells were transfected with a plasmid encoding the firefly luciferase gene under the control of the T7 promoter (pT7EMCLuc). All cells were transfected for 6 h at 37°C in OptiMEM (ThermoFisher Scientific) using the Lipofectamine 3000 system (Invitrogen, Carlsbad, CA, USA). A single cell suspension of target cells were added to effector cell monolayers and co-cultured in Ham’s F12 medium for 18 h at 37°C. Cells were lysed, and substrate was added to cell lysates (ProMega Luciferase Assay System) and immediately assayed for light output (fusion) using a BioTek Synergy Neo microplate luminometer. **(c)** (**i**) CHO-K1 effector cells transiently expressing wild type gB or gB Q181P and wild type gD, gH, and gL, and T7 polymerase were co-cultured with CHO-HVEM cells transiently expressing the luciferase gene. Luciferase-induced luminescence was quantitated as a measure of fusion. (**ii**) Detection of gB in the cell plasma membrane by CELISA. CHO-K1 cells were transfected with plasmids encoding HSV-1 gB or empty vector (EV) for 24 h. Cells were fixed with paraformaldehyde and then incubated with anti-gB monoclonal antibodies H126, H1359, and H1817. HRP-conjugated Protein A was added, followed by ABTS substrate. Absorbance was read at 405 nm. Wild type gB values were set to 100%. Results are the mean of three independent experiments. ns, not significant; Student’s *t-test*. **(d)** HSV-1 gB antibody binding sites. Space-filling rendering of prefusion and postfusion gB trimer with surface-exposed epitopes targeted by specific monoclonal antibodies. **(e)** Antigenic analysis of gB Q181P by dot blot. Lysates from CHO-K1 cells expressing gB Q181P or gB wild type were bound to nitrocellulose membrane then probed with a panel of antibodies targeting different domains (D) of gB. Fluorophore-conjugated secondary antibody was added for 20 min. Reactivity of the antibodies was quantitated relative to the wild type by densitometry (Azure Biosystems Image Studio). Results are the mean of three independent experiments. **, p ≤ 0.01; ***, p ≤ 0.001, ****, p ≤ 0.0001, Student’s t-test.

### Conclusions

gB undergoes dramatic structural changes from a meta-stable pre-fusion conformation to a stable post-fusion conformation during the HSV-cell membrane fusion process. The complicated structural dynamics and the complex molecular interactions are largely unclear but critically important. Here, we established and implemented a machine learning based approach, LINES, to evaluate the effects of gB fusion loop-MPR interactions on the stabilities of both pre- and post-fusion conformations. Based on which, a new and crucial interaction, Q181-R747, was identified. The molecular simulations with the Q181P mutation showed a reduced stability of the gB pre-fusion conformation. Meanwhile, experiments were also performed to test the effects of Q181P on fusion. Strikingly, the mutation of Q181 completely abrogated gB membrane fusion activity. The drastic reduction in cell-cell fusion upon Q181P mutation does confirm the importance of Q181-R747 interaction identified from our simulations. In addition, it suggests that – in light of the “loaded spring” pre-fusion conformation analogy – the fusion loop residue Q181 may be critically important in stabilizing the meta-stable pre-fusion conformation of gB. When this interaction is prevented via mutation or potentially disrupted by other molecules, the gB pre-fusion structure may be prematurely “unloaded” or “released”, leading to the final, stable post-fusion conformation. It is possible that the Q181-R747 interaction is modulated in the wild type gB by interactions with other glycoproteins such as gD, or gH/gL. Removal of this interaction via the Q181P mutation, may manually trigger gB without the need for the preceding glycoproteins involved in the viral fusion signaling cascade.

Inspection of the molecular simulation data reveals that the Q181P mutation disrupts the fusion loop secondary structures, weakens fusion loop-MPR residues interactions and destabilizes the pre-fusion conformation. It should be noted that there is a significant gap between stability of gB pre-fusion conformation (from machine learning and simulations) and cell-cell fusion reduction with Q181P mutant (from experiment). Our simulation data may provide a possible explanation but do not rule out other possibilities. For example, the disruption of the fusion loop from Q181P may disable the proper insertion of fusion loop into the membrane and lead to the fusion deficiency. Therefore, additional future experiments and simulations can be designed and conducted to fill the gap. Moving forward, it might be possible to inhibit the viral fusion process altogether by designing molecules or proteins to further strengthen the fusion-loop-MPR interaction. There is no HSV vaccine and no cure for latent infection. A stabilized form of pre-fusion gB may be an effective vaccine antigen, as has been the case for other viral fusion protein vaccines.^33^ It has been decades since there have been advances in antiviral drugs for HSV. The unclear mechanism of herpesviral fusion and entry has been a roadblock to developing drugs that inhibit entry. The results here may lay the groundwork for development of an HSV-specific fusion inhibitor, such as the clinically effective antiretroviral drugs for HIV. ^34, 35^

These results demonstrate how a union of simulation and experimental techniques can generate new leads when striving to understand the complex nature of biological interactions. Molecular simulations can capture the fine details of inter-atomic and inter-residue interactions, and the immense volume of information generated from the simulations can be processed with machine learning models like LINES that capture patterns in the data to understand primary interactions that occur. In this work, over 1000 inter-residue distances were captured during the MD simulations, but LINES was able to quickly evaluate and identify which fusion loop residues exhibited strong or weak interactions. The experimental techniques were then used to elucidate the impact of a single mutation on the entire molecular pathway and viral fusion as a whole. It was found that eliminating one strong pre-fusion interaction drastically altered the ability of gB to participate in viral membrane fusion, a finding that could have taken a very long time to discover if an unguided, trial-and-error approach to mutation had instead been used. Therefore, this approach could also be applied to theoretically any other biological system that can be simulated with MD simulations for which molecular interactions are important.

It should be noted that the diffusion model used in LINES can employ very flexible neural network frameworks, and the model is relatively simple to implement. Therefore, as the field of machine learning and unsupervised learning models continually improve, the neural networks used for LINES could also be updated to incorporate new developments without disrupting the overall LINES flow. One possible disadvantage in the current implementation of LINES is that reaction coordinates must be constructed from a linear combination of reaction coordinates for potentially non-linear reaction pathways, but this could potentially be avoided by merging the diffusion model with other reaction coordinate discovery techniques, such as Deep-TICA^36^. This combination of the gradient-based reaction coordinate discovery in LINES with the reaction coordinate flexibility of Deep-TICA could produce a very powerful tool for understanding molecular pathways.

## Materials and Methods

### System Setup

To generate a full-length pre-fusion gB structure, elements from the pre-fusion ecto-domain (PDBid: 6Z9M) and the post-fusion (PDBid: 5V2S) membrane region are used – shown in **Figure S1(a)** in Supporting Information. Cooper *et al*.^*25*^ noted that the transmembrane domain structures between pre- and post-fusion conformations should be similar, therefore we align the fusion loops (W174-Q181 and V259-Y265) from both conformations to superimpose the transmembrane domain onto the pre-fusion structure. Missing residues are filled in with SWISS modeler^37^, producing the full-length pre-fusion gB structure in **Figure S1(b)** in Supporting Information.

The post-fusion gB structure uses the full-length structure (PDBid: 5V2S) but then truncates domains II-IV to reduce the size of the simulation box, as shown in **Figure S1(c)** in Supporting Information. Missing residues in the transmembrane region are also filled in with SWISS modeler.^37^ We have simulated a full-length post-fusion structure for a short time period, but no discernible difference of the domain I/MPR/transmembrane domain/CTD was observed over 10 ns simulations when compared to the truncated structure. Therefore, the truncated structure is adopted, allowing for longer MD simulations to be performed. Since this work focuses on the domain I-MPR interactions, the truncated II-IV domains should not impact the results reported here.

After obtaining pre-fusion and post-fusion gB conformations, the remaining steps for setting up the simulation box are identical. For both gB conformations, the protein is centered in the simulation box of size 15 *nm* × 15 *nm* × 22 *nm* and is embedded in a lipid membrane (2 POPE/1 POPC/1 DOPC/1 Cholesterol ratio^38^) across the x/y plane (15 *nm* × 15 *nm*). The system is then solvated with TIP3P^39^ molecules. The MD simulations are simulated in GROMACS 2020.4^40^ that has been patched with PLUMED 2.8.^41^ The CHARMM36 forcefield^42^ is adopted and the inputs to the GROMACS software are generated with CHARMM-GUI.^43^ The system is minimized and then relaxed at the NVT and NPT ensembles for 1 ns each, with the pressure set to 1 bar and the temperature set to 310K.

### MD Simulations

After setting up the systems, MD simulations were performed and a set of molecular coordinates were saved during the simulations for ML model. Molecular coordinates (or descriptors), such as atomic distances, bond angles, or dihedral angles that contain molecular reference groups – were fed into the ML model, so that we can analyze and predict the important interactions among gB residues and lipid membranes. The molecular coordinates used in this work, as illustrated in **Fig. 2(a)**, are defined from the distances between the fusion loop residues (W174-Q181, V259-Y265) and the 10 closest MPR residues in the starting pre- and post-fusion structures. Since the 10 closest residues to a fusion loop may be different from different monomers of the gB trimer or whether the system is in the pre- or post-fusion conformation, we select a set of distances so that each fusion loop residue has a distance measured with the same MPR residues regardless of chain or pre/post-fusion conformation. This will allow for a comparison between the pre-fusion and post-fusion conformations and also provide three estimates (from the three monomers) of the fusion loop-MPR interactions. An additional set of molecular coordinates will also be defined to capture the interactions between fusion loop residues with the lipid membrane: the closest distance to each type of lipid will be saved for each fusion loop residue.

### Machine Learning

We implemented our machine learning model, LINES^26, 27^, to identify reaction pathways and strong interactions in protein systems. Typically, this would be accomplished through iterations of biased MD simulation and ML to iteratively improve reaction coordinates and sample more conformations. As this work focuses on identifying the fusion loop-MPR interactions that stabilize the pre- and post-fusion conformations instead of simulating the reaction in its entirety, the unbiased MD simulations described in the previous section should be sufficient to observe interactions already in place.

The ML model will learn gradients of the free energy surface (FES), allowing for optimization techniques such as time-lagged independent component analysis (TICA)^44^ to identify the slow degrees of freedom in the system. To capture the FES gradients, a diffusion model – also sometimes referred to as score-based generative models^45^ – will be used. Diffusion models^30, 31^ learn gradients of the log-probability distribution. Since the log-probability is proportional to the free energy of the system, a diffusion model also learns the FES gradients. In score-based generative models, the input data – the molecular coordinates from MD simulations – will have gradual Gaussian noise added through a sequence of intensifying Gaussian noise.

A neural network is then trained to try “denoising” the data by estimating the Stein score. For a data distribution *q*(*x*_0_) and a sequence of noise levels 0 < σ_1_ < σ_2_ < ⋯ < σ_*t*_, a data point can be perturbed from *x*_0_ to *x*_*t*_ with a Gaussian noise distribution:

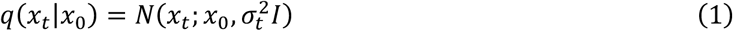

The deep neural network *s*_*θ*_(*x, t*) tries to predict the Stein score function ∇_*x*_ log *q*(*x*_*t*_). We should note the subscript *t* here does not refer to the simulation time but a time along a latent distribution for a stochastic differential equation.^45^ The Stein score is a function of the data itself and is a vector field pointing to directions along which the probability density function will grow the fastest. In the context of FES gradients, the Stein score models the forces between molecular reference groups in the protein system. This score function has a succinct form^30, 46^ to provide the following loss function for model training:

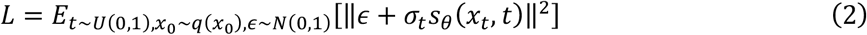

In this equation, *U*(0,1) and *N*(0,1) represent the uniform and normal distribution from 0 to 1. The noisy data is computed as *x*_*t*_ = *µ*_*t*_*x*_0_ + *σ*_*t*_*ϵ. µ*_*t*_ and *σ*_*t*_ represent the mean drifts and mean deviations for a stochastic process up to a specified time *t*, and *ϵ* represents noise from a multivariate independent Gaussian distribution. The drifts and deviations are estimated using the sub-variance preserving stochastic differential equation form described by Yang *et al*.^45^: 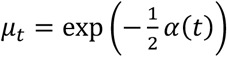 and *σ*_*t*_ = 1 − exp(−*α*(*t*)), where 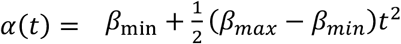. The conventional values of *β*_*min*_ = 0.001 and *β*_*max*_ = 3 are used for this work.

One benefit of using diffusion models is that the neural network architecture can be very loose as long as the output dimensionality equals the input dimensionality. Therefore, if there are *N*_*MC*_ molecular coordinates inputted to the model, the neural network will output a vector with *N*_*MC*_ components. The neural network uses a U-shaped network with 6 dense residual layers of 256-128-64-64-128-256 hidden units. The residual network skips connections to prevent gradient vanishing while training the neural network. Hidden layers in the model use a Swish activation function (*f*(*x*) = *x* ∗ *Sigmoid*(*x*)) and the output layer has a linear activation function. Before inputting the molecular coordinates into the model, the data is normalized to have mean zero and a standard deviation of 1. The neural network weights are initialized according to the Glorot and Bengio scheme^47^ and utilize L_2_-regularization with a coefficient set to 5 × 10^−5^.

The ADAM^48^ optimizer is used to train the neural network with a learning rate of 1 × 10^−4^. A batch size of 1000 is used, and the training is terminated after 1000 epochs. However, if the loss function does not decrease for more than 100 epochs, training is terminated to prevent overfitting.

### Reaction Coordinate Prediction

In this work, the diffusion model is leveraged to estimate FES gradients. Gradients are useful when trying to isolate important interactions, because the gradients can provide information about the reaction pathways before the entire reaction is sampled.^49^ The neural network *s*_*θ*_(*x*_*t*_, *t*) will be evaluated for each data sample at a “small” time in the stochastic diffusion process (e.g., *t* = 10^−5^) to produce the FES gradient estimates. The gradients are then used to estimate a covariance matrix between samples separated by a given number of lagged simulation timesteps Δ*l*:

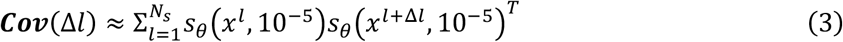

The important reaction coordinates are computed via eigenvalue decomposition of the gradient-based autocorrelation equation inspired by time-lagged independent component analysis (TICA)^44^:

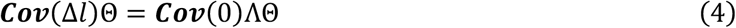

In this equation, Θ is a coefficient matrix, where the eigenvectors are sorted so that the corresponding eigenvalues Λ are in descending order. As the covariance matrix is not guaranteed to produce non-imaginary eigenvalues, ***Cov***(Δ*l*) is averaged with its transpose. The reaction coordinate is taken from the first eigenvector of Θ. Since several hundred molecular coordinates are inputted into the ML model, which makes analysis of the reaction coordinate very difficult. To overcome this, we compute a separate reaction coordinate for each fusion loop residue. The molecular coordinate coefficients *θ*_*i*_ for each reaction coordinate are extracted as the components of the eigenvector with the leading eigenvalue, and the reaction coordinate is computed as the weighted summation of each molecular coordinate *d*_*i*_, as *RC* = ∑_*i*_ *θ*_*i*_ *d*_*i*_.

### Interaction Strength and Coordination of Correlations

After obtaining the reaction coordinate for every fusion loop residue, we evaluated the interaction strength from the given fusion loop residue by examining whether the corresponding reaction coordinate captures the coordinated motions or reactions. A strength metric, Coordination of Correlations (CC), is defined based on the weighted L2-norm on the covariance between a reaction coordinate and each of its associated molecular coordinates^50^:

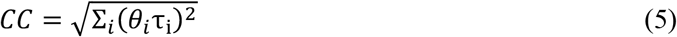

Here *τ*_*i*_ is the Kendall-Tau correlation^51^ between the reaction coordinate and molecular coordinate *i*. The Kendall-Tau correlation is computed from a set of (*x*_1_, *y*_1_), …, (*x*_*n*_, *y*_*n*_) joint random variables and measures the number of concordant pairs of observations with the following equation:

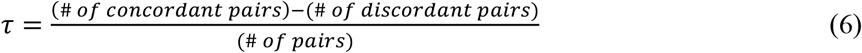

The Kendall-Tau correlation is used to measure how strong of a link exists between two variables (i.e., the reaction coordinate and a molecular coordinate). If there are several uncorrelated molecular coordinates used in a single reaction coordinate, then the reaction coordinate will not have a strong correlation with any individual molecular coordinate, even though it is composed of a linear combination of said molecular coordinates.

This “strength” metric is computed for each individual reaction coordinate and is bounded between values of 0 and 1 if the molecular coordinate coefficients are rescaled to have a norm of 1. This metric groups all of the collective motions of the molecular coordinates with the interaction strength predicted by the ML model. For example, a stability of 0 indicates that every molecular coordinate is essentially random noise and has no coordinated motion, so the interactions involved with the fusion loop residue are not very strong. Alternatively, a stability of 1 indicates that all of the molecular coordinates move in a coordinated motion that can be entirely captured with the reaction coordinate.

### Cells

Chinese hamster ovary (CHO-K1) cells (American Type Culture Collection (ATCC), Manassas, VA, USA) were propagated in Ham’s F12 nutrient mixture (Gibco/Life Technologies, Grand Island, NY, USA) supplemented with 10% fetal bovine serum (FBS) (Atlanta Biologicals, Atlanta, GA, USA). CHO-K1 cells expressing the human herpesvirus entry mediator (CHO-HVEM or M1A cells) ^52^ were obtained from G. Cohen and R. Eisenberg, University of Pennsylvania. CHO-HVEM cells were propagated in Ham’s F12 nutrient mixture supplemented with 10% FBS, 150 µg/mL puromycin (Sigma Aldrich, St. Louis - MO, USA), and 250 µg/mL G418 sulfate (Sigma Aldrich, St. Louis, MO, USA). Cells were subcultured in non-selective medium for at least two passages before use in experiments.

### Construction of HSV-1 gB with a point mutation (Q181P) in domain I

The wild type HSV-1 strain KOS gB sequence^53^ was synthesized with the *NsiI* and *EcoRI* flanking sites upstream and downstream of gB, respectively, using GeneArt (ThermoFisher Scientific (Regensburg, Germany) and inserted into a pMA vector (ThermoFisher Scientific) using the Q5 Site-Directed Mutagenesis Kit (New England Biolabs, Ipswich, MA, USA) and mutagenic primers, a single nucleotide point mutation from glutamine to proline at residue 181 was introduced in the gB sequence. The primer pairs consisted of a sense primer 5′-CCG CTA CTC CCC GTT TAT GGG GA-3′ and an antisense primer 5′-TGG CCG AAC CAC ACC-3′. The generated plasmid construct was digested with *NsiI, Ase I*, and *EcoRI* enzymes, while the pCAGGS/MCS^54, 55^ backbone was digested with *NsiI* and *EcoRI* enzymes. The digestion products were gel purified with a QiaQuick Gel Extraction Kit (Qiagen). The gB gene fragment and pCAGGS/MCS DNAs were then ligated using Instant Sticky-End Ligase (New England Biolabs) and transformed into competent *E. coli* DH5-α cells (New England Biolabs), generating pAM26 (gB Q181P). All restriction enzymes were from New England Biolabs. All plasmids were sequence-verified.

### Antibodies to gB

Anti-HSV-1 gB mouse monoclonal antibodies H126 ^56, 57^, H1359 ^58^, and H1817 ^58^ were purchased from Virusys, Taneytown, MD, USA. Anti-gB monoclonal antibodies DL16 (oligomer specific), SS10, and SS106, and rabbit polyclonal serum to gB R68 were provided by G. Cohen and R. Eisenberg, University of Pennsylvania.^59, 60^

### Analysis of cell surface-expressed gB by CELISA

CHO-K1 cells in 96-well plates were transfected with plasmids encoding gB Q181P, gB wild type, or empty plasmid vector with Lipofectamine 3000 (Invitrogen, Carlsbad, CA, USA). Cells were cultured for 18 h and then fixed in 4% paraformaldehyde. Fixed cells were blocked with 3% BSA in PBS for 2 h. Anti-gB monoclonal antibodies H126, H1359, and H1817 in 3% BSA in PBS were added overnight at 4°C. Protein A conjugated to horseradish peroxidase (Invitrogen) was added for 2 h at room temperature. Substrate 2,2′-Azinobis [3-ethylbenzothiazoline-6-sulfonic acid]-diammonium salt (ABTS; Thermo Fisher Scientific) was added, and absorbance was measured at 405 nm using a BioTek microplate reader.

### Luciferase reporter assay for cell-cell fusion

CHO-K1 (effector) cells were transfected with plasmids encoding T7 RNA polymerase (pCAGT7), HSV-1 (strain KOS) plasmids pPEP98 (wild type gB) or pAM26 (gB Q181P), pPEP99 (wild type gD), pPEP100 (wild type gH), and pPEP101 (wild type gL) ^61^. CHO-HVEM (target) cells were transfected with a plasmid encoding the firefly luciferase gene under the control of the T7 promoter (pT7EMCLuc). All cells were transfected for 6 h at 37°C in OptiMEM (ThermoFisher Scientific) using the Lipofectamine 3000 system (Invitrogen, Carlsbad, CA, USA). A single cell suspension of target cells were added to effector cell monolayers and co-cultured in Ham’s F12 medium for 18 h at 37°C. Cells were lysed using the ProMega Luciferase Assay System, and lysates were frozen and thawed. Substrate was added to cell lysates and immediately assayed for light output (luciferase activity; fusion) using a BioTek Synergy Neo microplate luminometer.

### Immuno dotblot analysis of HSV-1 gB

Transfected CHO-K1 cell lysates were diluted two-fold in serum-free, bicarbonate-free DMEM with 0.2% bovine serum albumin (BSA) and 5 mM (each) HEPES, MES (morpholineethanesulfonic acid), and sodium succinate. Approximately 10^7^ genome copies of HSV-1 were blotted directly to a nitrocellulose membrane using a Minifold dot blot system (Whatman). Membranes were blocked and then incubated with antibodies to gB. After incubation with fluorescent-conjugated secondary antibody for 20 min, images were obtained with an Azure Biosystems imager. Densitometry was performed with Image Studio. Wild type gB reactivity with each antibody was normalized to 1.00.

## Author contributions

R.E.O. developed the methods and performed the simulations. A.O.M. and M.A.H performed experiments. P.D., A.V.N. and J.L. designed and guided the research. All authors contributed to the interpretation of the data and writing/review of the manuscript.

## Data availability

Inputs needed to run the simulations and instructions of how to employ these files in PLUMED are available at GitHub at https://github.com/reodstrcil/gB. All the simulations are performed using open-source software as described in the Materials and Methods sections.

## Acknowledgements

Research reported in this publication was supported by the National Institute of General Medical Sciences of the National Institutes of Health under award numbers R01GM152745 and T32GM008336. Computational resources were provided by Expanse CPU at SDSC through allocation MCB170012 from the Advanced Cyberinfrastructure Coordination Ecosystem: Services & Support (ACCESS) program, which is supported by National Science Foundation grants #2138259, #2138286, #2138307, #2137603, and #2138296.

