## Supplemental Materials for "Modulation of specific interactions within a viral fusion protein predicted from machine learning blocks membrane fusion"

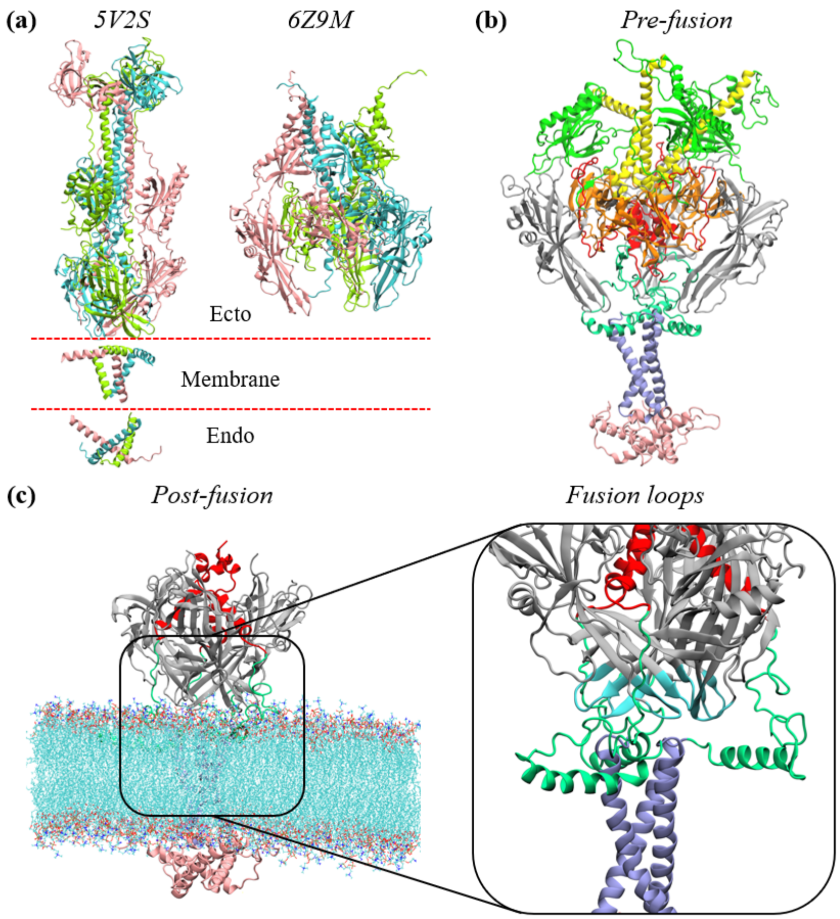


**Figure S1**: Preparation of the gB pre/post-fusion systems. **(a)** The structures for gB in post-fusion (PDBid: 5V2S) and pre-fusion (PDBid: 6Z9M) are available. Different colors represent the 3 monomers of the gB trimer. The transmembrane and endo-domain portions of gB are only available for the post-fusion structure, so the full-length pre-fusion structure is generated by superimposing the membrane and endo-domain regions of the post-fusion structure onto the pre-fusion structure. **(b)** The pre-fusion structure of gB is shown and are colored based on the domains: DI (gray), DII (light green), DIII (yellow), DIV (orange), DV (red), membrane proximal region (MPR) (turquoise), transmembrane domain (purple), and cytoplasmic tail domain (pink). **(c)** An example of how the fusion loops (blue) in the truncated post-fusion structure are embedded in the membrane is shown. The truncated post-fusion structure has regions DII-DIV removed.

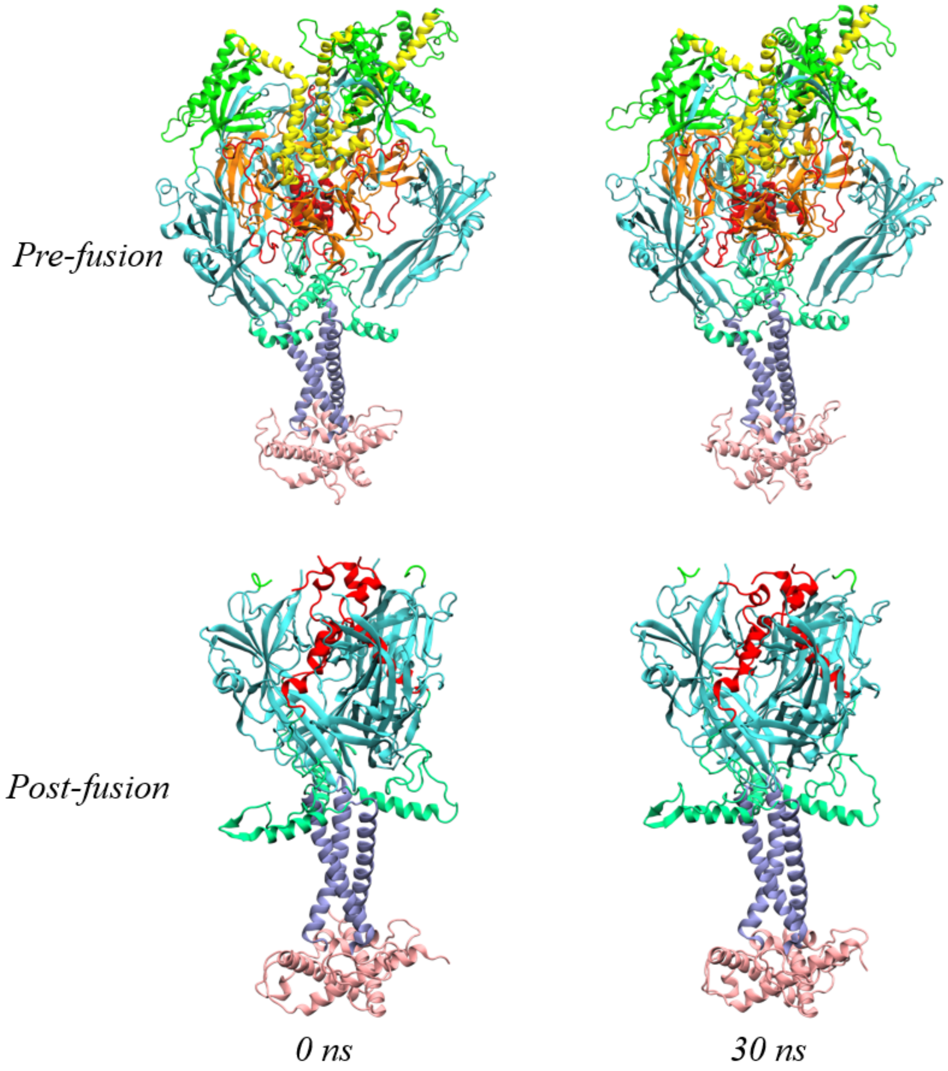


**Figure S2**: gB conformations over 30 ns trajectory. A comparison of the starting conformation at 0 ns and the ending conformation at 30 ns for both the pre-fusion and post-fusion systems reveals stable binding poses. The different domains are colored as follows: cyan for domain I, green for domain II, yellow for domain III, orange for domain IV, red for domain V, light green for membrane-proximal region, purple for transmembrane domain, and pink for cytoplasm tail domain.

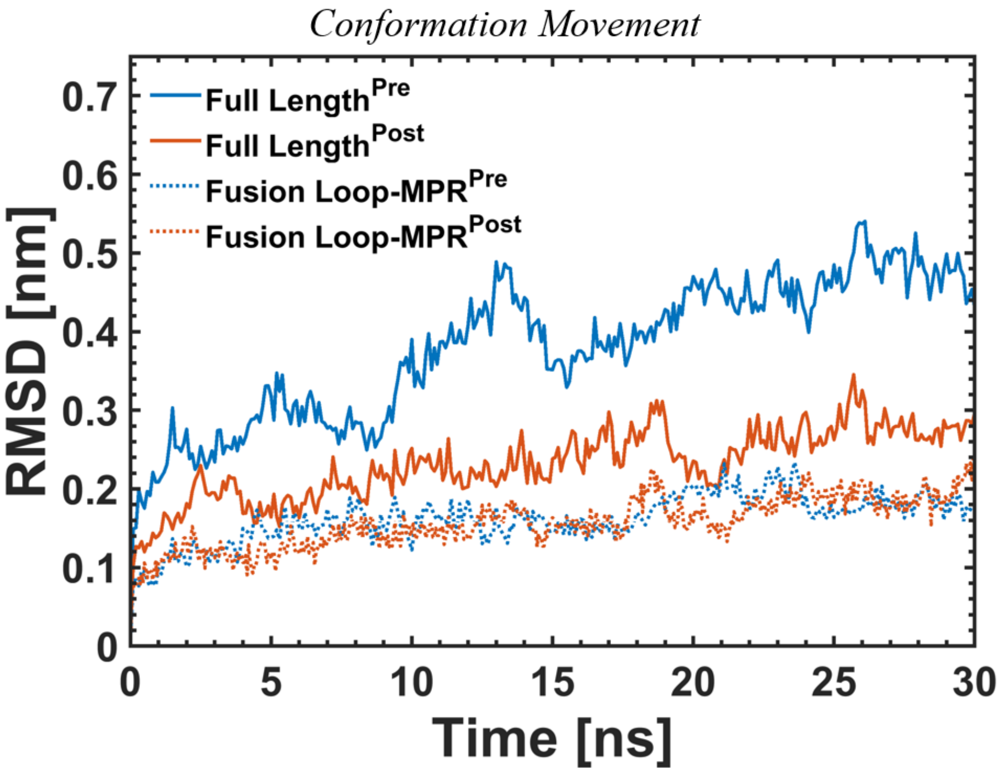


**Figure S3**: Conformational Movement of the Pre- and Post-fusion Systems. The root mean squared deviation (RMSD) for the full-length and fusion loop/membrane proximal region (MPR) pre- and post-fusion gB structures is shown. The post-fusion gB structure is stable throughout 30 ns of MD simulation, while the full-length pre-fusion gB structure shows motion. However, this motion is due to domain 3 of the pre-fusion structure, which does not interact in close proximity with the fusion loops or the MPR.

### Pre-fusion Reaction Coordinate Coefficients & Coordinate Correlations

*Fusion Loop Residue*

*MPR Residue*

| | TRP174 | PHE175 | GLY176 | HSD177 | ARG178 | TYR179 | SER180 | GLN181 | VAL259 | GLU260 | ALA261 | PHE262 | HSD263 | ARG264 | TYR265 |
|---|---|---|---|---|---|---|---|---|---|---|---|---|---|---|---|
| THR721 | 0.000 | 0.000 | 0.000 | 0.000 | 0.000 | 0.000 | 0.467 | 0.000 | 0.000 | 0.000 | 0.000 | 0.000 | 0.000 | 0.000 | 0.000 |
| VAL722 | 0.000 | 0.000 | 0.000 | 0.000 | 0.000 | 0.148 | 0.287 | 0.398 | 0.000 | 0.000 | 0.000 | 0.000 | 0.000 | 0.000 | 0.000 |
| ILE723 | 0.000 | 0.184 | 0.000 | 0.000 | 0.000 | 0.148 | 0.158 | 0.052 | 0.000 | 0.000 | 0.000 | 0.000 | 0.000 | 0.181 | 0.108 |
| HSD724 | 0.106 | 0.106 | 0.053 | 0.040 | 0.049 | 0.148 | 0.144 | 0.305 | 0.000 | 0.000 | 0.000 | 0.000 | 0.000 | 0.000 | 0.000 |
| ALA725 | 0.092 | 0.172 | 0.165 | 0.000 | 0.210 | 0.265 | 0.139 | 0.265 | 0.131 | 0.000 | 0.000 | 0.000 | 0.000 | 0.088 | 0.184 |
| ASP726 | 0.103 | 0.103 | 0.267 | 0.106 | 0.143 | 0.191 | 0.199 | 0.240 | 0.125 | 0.027 | 0.000 | 0.000 | 0.312 | 0.165 | 0.246 |
| ALA727 | 0.061 | 0.086 | 0.084 | 0.397 | 0.146 | 0.199 | 0.123 | 0.095 | 0.134 | 0.085 | 0.000 | 0.000 | 0.000 | 0.238 | 0.000 |
| ASN728 | 0.064 | 0.010 | 0.181 | 0.297 | 0.114 | 0.139 | 0.029 | 0.098 | 0.152 | 0.019 | 0.199 | 0.000 | 0.000 | 0.154 | 0.009 |
| ALA729 | 0.000 | 0.053 | 0.196 | 0.206 | 0.159 | 0.264 | 0.067 | 0.049 | 0.353 | 0.015 | 0.000 | 0.000 | 0.000 | 0.378 | 0.070 |
| ALA730 | 0.000 | 0.220 | 0.062 | 0.396 | 0.175 | 0.028 | 0.020 | 0.228 | 0.256 | 0.000 | 0.000 | 0.000 | 0.000 | 0.158 | 0.012 |
| MET731 | 0.136 | 0.285 | 0.267 | 0.025 | 0.117 | 0.039 | 0.059 | 0.183 | 0.000 | 0.000 | 0.000 | 0.000 | 0.286 | 0.251 | 0.001 |
| PHE732 | 0.000 | 0.000 | 0.000 | 0.000 | 0.226 | 0.220 | 0.064 | 0.000 | 0.000 | 0.000 | 0.000 | 0.000 | 0.000 | 0.000 | 0.000 |
| ALA733 | 0.000 | 0.098 | 0.039 | 0.348 | 0.343 | 0.257 | 0.227 | 0.267 | 0.000 | 0.000 | 0.000 | 0.000 | 0.000 | 0.000 | 0.000 |
| GLY734 | 0.000 | 0.000 | 0.000 | 0.066 | 0.242 | 0.000 | 0.000 | 0.000 | 0.000 | 0.000 | 0.000 | 0.000 | 0.000 | 0.000 | 0.000 |
| LEU735 | 0.000 | 0.000 | 0.000 | 0.000 | 0.276 | 0.000 | 0.000 | 0.000 | 0.000 | 0.000 | 0.000 | 0.000 | 0.000 | 0.000 | 0.000 |
| ASP744 | 0.223 | 0.177 | 0.149 | 0.025 | 0.100 | 0.146 | 0.149 | 0.194 | 0.000 | 0.000 | 0.000 | 0.000 | 0.000 | 0.000 | 0.099 |
| LEU745 | 0.243 | 0.140 | 0.331 | 0.050 | 0.102 | 0.145 | 0.175 | 0.113 | 0.000 | 0.000 | 0.000 | 0.000 | 0.398 | 0.203 | 0.302 |
| GLY746 | 0.081 | 0.089 | 0.220 | 0.199 | 0.127 | 0.194 | 0.157 | 0.156 | 0.426 | 0.122 | 0.132 | 0.164 | 0.131 | 0.205 | 0.157 |
| ARG747 | 0.242 | 0.238 | 0.081 | 0.000 | 0.000 | 0.270 | 0.220 | 0.113 | 0.070 | 0.000 | 0.189 | 0.115 | 0.116 | 0.256 | 0.095 |
| ALA748 | 0.210 | 0.214 | 0.081 | 0.000 | 0.504 | 0.256 | 0.205 | 0.135 | 0.102 | 0.000 | 0.119 | 0.034 | 0.091 | 0.285 | 0.177 |
| VAL749 | 0.301 | 0.000 | 0.121 | 0.000 | 0.000 | 0.221 | 0.283 | 0.185 | 0.000 | 0.000 | 0.174 | 0.000 | 0.215 | 0.080 | 0.183 |
| GLY750 | 0.216 | 0.330 | 0.008 | 0.000 | 0.000 | 0.240 | 0.095 | 0.155 | 0.232 | 0.179 | 0.166 | 0.149 | 0.232 | 0.081 | 0.201 |
| LYS751 | 0.197 | 0.342 | 0.467 | 0.000 | 0.000 | 0.000 | 0.026 | 0.130 | 0.013 | 0.117 | 0.386 | 0.137 | 0.157 | 0.246 | 0.201 |
| VAL752 | 0.172 | 0.304 | 0.185 | 0.000 | 0.000 | 0.000 | 0.181 | 0.136 | 0.148 | 0.074 | 0.203 | 0.083 | 0.224 | 0.161 | 0.200 |
| VAL753 | 0.000 | 0.000 | 0.000 | 0.000 | 0.000 | 0.000 | 0.000 | 0.000 | 0.067 | 0.164 | 0.205 | 0.219 | 0.322 | 0.009 | 0.277 |
| MET754 | 0.000 | 0.283 | 0.397 | 0.090 | 0.000 | 0.000 | 0.000 | 0.059 | 0.242 | 0.072 | 0.044 | 0.178 | 0.087 | 0.103 | 0.251 |
| GLY755 | 0.326 | 0.121 | 0.164 | 0.142 | 0.000 | 0.145 | 0.208 | 0.154 | 0.295 | 0.192 | 0.271 | 0.204 | 0.147 | 0.077 | 0.308 |
| ILE756 | 0.078 | 0.140 | 0.070 | 0.075 | 0.000 | 0.000 | 0.163 | 0.000 | 0.080 | 0.078 | 0.254 | 0.215 | 0.070 | 0.213 | 0.228 |
| VAL757 | 0.000 | 0.000 | 0.000 | 0.093 | 0.000 | 0.000 | 0.000 | 0.000 | 0.194 | 0.167 | 0.313 | 0.000 | 0.000 | 0.000 | 0.000 |
| GLY758 | 0.073 | 0.104 | 0.063 | 0.120 | 0.136 | 0.115 | 0.132 | 0.000 | 0.197 | 0.236 | 0.144 | 0.300 | 0.302 | 0.236 | 0.199 |
| GLY759 | 0.065 | 0.168 | 0.189 | 0.134 | 0.273 | 0.251 | 0.049 | 0.143 | 0.180 | 0.153 | 0.175 | 0.214 | 0.003 | 0.131 | 0.126 |
| VAL760 | 0.000 | 0.000 | 0.000 | 0.142 | 0.078 | 0.000 | 0.000 | 0.000 | 0.074 | 0.026 | 0.054 | 0.000 | 0.000 | 0.000 | 0.000 |
| VAL761 | 0.000 | 0.000 | 0.016 | 0.157 | 0.369 | 0.000 | 0.000 | 0.000 | 0.140 | 0.081 | 0.073 | 0.000 | 0.000 | 0.000 | 0.000 |
| SER762 | 0.186 | 0.087 | 0.134 | 0.198 | 0.149 | 0.149 | 0.007 | 0.062 | 0.124 | 0.102 | 0.051 | 0.188 | 0.105 | 0.180 | 0.000 |
| ALA763 | 0.000 | 0.283 | 0.281 | 0.172 | 0.166 | 0.323 | 0.179 | 0.000 | 0.097 | 0.159 | 0.224 | 0.000 | 0.000 | 0.000 | 0.000 |
| VAL764 | 0.000 | 0.000 | 0.000 | 0.200 | 0.000 | 0.000 | 0.000 | 0.000 | 0.000 | 0.071 | 0.000 | 0.000 | 0.000 | 0.000 | 0.000 |
| SER765 | 0.135 | 0.014 | 0.174 | 0.155 | 0.263 | 0.207 | 0.116 | 0.183 | 0.120 | 0.107 | 0.186 | 0.144 | 0.185 | 0.162 | 0.000 |
| GLY766 | 0.155 | 0.161 | 0.203 | 0.157 | 0.196 | 0.160 | 0.239 | 0.104 | 0.220 | 0.067 | 0.277 | 0.136 | 0.207 | 0.095 | 0.000 |
| VAL767 | 0.000 | 0.000 | 0.177 | 0.231 | 0.099 | 0.000 | 0.000 | 0.000 | 0.000 | 0.172 | 0.148 | 0.107 | 0.000 | 0.000 | 0.000 |
| SER768 | 0.000 | 0.000 | 0.000 | 0.000 | 0.148 | 0.000 | 0.000 | 0.000 | 0.160 | 0.024 | 0.124 | 0.189 | 0.123 | 0.000 | 0.000 |
| SER769 | 0.112 | 0.100 | 0.055 | 0.206 | 0.179 | 0.097 | 0.068 | 0.086 | 0.181 | 0.165 | 0.219 | 0.112 | 0.202 | 0.284 | 0.096 |
| PHE770 | 0.120 | 0.000 | 0.127 | 0.000 | 0.000 | 0.187 | 0.000 | 0.000 | 0.000 | 0.174 | 0.157 | 0.228 | 0.100 | 0.000 | 0.068 |
| MET771 | 0.179 | 0.000 | 0.183 | 0.177 | 0.142 | 0.167 | 0.074 | 0.200 | 0.000 | 0.098 | 0.129 | 0.151 | 0.108 | 0.094 | 0.117 |
| SER772 | 0.248 | 0.000 | 0.000 | 0.000 | 0.061 | 0.000 | 0.000 | 0.000 | 0.067 | 0.000 | 0.149 | 0.077 | 0.082 | 0.146 | 0.155 |
| ASN773 | 0.000 | 0.000 | 0.000 | 0.000 | 0.000 | 0.000 | 0.000 | 0.000 | 0.252 | 0.000 | 0.000 | 0.212 | 0.000 | 0.160 | 0.063 |
| PHE775 | 0.209 | 0.000 | 0.000 | 0.039 | 0.176 | 0.185 | 0.000 | 0.000 | 0.000 | 0.000 | 0.227 | 0.000 | 0.000 | 0.000 | 0.130 |
| CC | 0.291 | 0.342 | 0.291 | 0.380 | 0.318 | 0.261 | 0.299 | 0.415 | 0.425 | 0.386 | 0.410 | 0.454 | 0.424 | 0.362 | 0.378 |

**Figure S4**: Reaction Coordinate Coefficients and Stabilities for Prefusion gB. The coefficients between the fusion loop residues (columns) and the membrane proximal region (MPR) residues (rows) are colored on a green scale to denote the magnitude of the coefficients. The bottom row shows the "stability" of each fusion loop residue, which is a measure of how well the reaction coordinate captures a coordinated motion of MPR residues.

#### Post-fusion Reaction Coordinate Coefficients & Coordinate Correlations

*Fusion Loop Residue* (columns); *MPR Residue* (rows)

| | TRP174 | PHE175 | GLY176 | HSD177 | ARG178 | TYR179 | SER180 | GLN181 | VAL259 | GLU260 | ALA261 | PHE262 | HSD263 | ARG264 | TYR265 |
|---|---|---|---|---|---|---|---|---|---|---|---|---|---|---|---|
| THR721 | 0.000 | 0.000 | 0.000 | 0.000 | 0.000 | 0.000 | 0.209 | 0.000 | 0.000 | 0.000 | 0.000 | 0.000 | 0.000 | 0.000 | 0.000 |
| VAL722 | 0.000 | 0.000 | 0.000 | 0.000 | 0.000 | 0.043 | 0.194 | 0.247 | 0.000 | 0.000 | 0.000 | 0.000 | 0.000 | 0.000 | 0.000 |
| ILE723 | 0.000 | 0.251 | 0.000 | 0.000 | 0.000 | 0.082 | 0.221 | 0.152 | 0.000 | 0.000 | 0.000 | 0.000 | 0.000 | 0.102 | 0.080 |
| HSD724 | 0.151 | 0.114 | 0.277 | 0.003 | 0.245 | 0.163 | 0.285 | 0.179 | 0.000 | 0.000 | 0.000 | 0.000 | 0.000 | 0.000 | 0.000 |
| ALA725 | 0.212 | 0.118 | 0.098 | 0.000 | 0.145 | 0.140 | 0.145 | 0.068 | 0.224 | 0.000 | 0.000 | 0.000 | 0.000 | 0.208 | 0.091 |
| ASP726 | 0.194 | 0.209 | 0.081 | 0.045 | 0.339 | 0.117 | 0.106 | 0.100 | 0.145 | 0.065 | 0.000 | 0.000 | 0.070 | 0.183 | 0.059 |
| ALA727 | 0.222 | 0.131 | 0.212 | 0.052 | 0.173 | 0.337 | 0.187 | 0.147 | 0.060 | 0.365 | 0.000 | 0.000 | 0.000 | 0.148 | 0.000 |
| ASN728 | 0.049 | 0.142 | 0.191 | 0.015 | 0.289 | 0.016 | 0.140 | 0.204 | 0.201 | 0.155 | 0.142 | 0.000 | 0.000 | 0.263 | 0.245 |
| ALA729 | 0.000 | 0.137 | 0.116 | 0.159 | 0.242 | 0.192 | 0.301 | 0.132 | 0.050 | 0.255 | 0.000 | 0.000 | 0.000 | 0.196 | 0.324 |
| ALA730 | 0.000 | 0.053 | 0.231 | 0.306 | 0.192 | 0.266 | 0.278 | 0.219 | 0.154 | 0.000 | 0.000 | 0.000 | 0.000 | 0.236 | 0.152 |
| MET731 | 0.142 | 0.179 | 0.067 | 0.079 | 0.033 | 0.209 | 0.066 | 0.272 | 0.000 | 0.000 | 0.000 | 0.000 | 0.389 | 0.282 | 0.190 |
| PHE732 | 0.000 | 0.000 | 0.000 | 0.000 | 0.301 | 0.093 | 0.070 | 0.000 | 0.000 | 0.000 | 0.000 | 0.000 | 0.000 | 0.000 | 0.000 |
| ALA733 | 0.000 | 0.510 | 0.365 | 0.230 | 0.177 | 0.026 | 0.157 | 0.027 | 0.000 | 0.000 | 0.000 | 0.000 | 0.000 | 0.000 | 0.000 |
| GLY734 | 0.000 | 0.000 | 0.000 | 0.233 | 0.134 | 0.000 | 0.000 | 0.000 | 0.000 | 0.000 | 0.000 | 0.000 | 0.000 | 0.000 | 0.000 |
| LEU735 | 0.000 | 0.000 | 0.000 | 0.000 | 0.090 | 0.000 | 0.000 | 0.000 | 0.000 | 0.000 | 0.000 | 0.000 | 0.000 | 0.000 | 0.000 |
| ASP744 | 0.159 | 0.127 | 0.298 | 0.053 | 0.220 | 0.169 | 0.121 | 0.095 | 0.000 | 0.000 | 0.000 | 0.000 | 0.000 | 0.000 | 0.197 |
| LEU745 | 0.076 | 0.151 | 0.330 | 0.040 | 0.262 | 0.229 | 0.225 | 0.219 | 0.000 | 0.000 | 0.000 | 0.000 | 0.128 | 0.258 | 0.088 |
| GLY746 | 0.186 | 0.163 | 0.089 | 0.344 | 0.102 | 0.180 | 0.124 | 0.252 | 0.285 | 0.059 | 0.117 | 0.178 | 0.123 | 0.223 | 0.082 |
| ARG747 | 0.156 | 0.033 | 0.244 | 0.000 | 0.000 | 0.143 | 0.224 | 0.133 | 0.195 | 0.000 | 0.126 | 0.112 | 0.232 | 0.186 | 0.083 |
| ALA748 | 0.078 | 0.211 | 0.178 | 0.000 | 0.027 | 0.070 | 0.159 | 0.108 | 0.122 | 0.000 | 0.171 | 0.317 | 0.187 | 0.041 | 0.026 |
| VAL749 | 0.107 | 0.000 | 0.166 | 0.000 | 0.000 | 0.114 | 0.186 | 0.139 | 0.000 | 0.000 | 0.061 | 0.000 | 0.200 | 0.001 | 0.053 |
| GLY750 | 0.109 | 0.194 | 0.266 | 0.000 | 0.000 | 0.090 | 0.263 | 0.177 | 0.217 | 0.219 | 0.239 | 0.061 | 0.292 | 0.341 | 0.034 |
| LYS751 | 0.508 | 0.271 | 0.472 | 0.000 | 0.000 | 0.000 | 0.139 | 0.147 | 0.296 | 0.114 | 0.031 | 0.195 | 0.102 | 0.123 | 0.089 |
| VAL752 | 0.222 | 0.151 | 0.108 | 0.000 | 0.000 | 0.000 | 0.044 | 0.153 | 0.229 | 0.322 | 0.298 | 0.217 | 0.204 | 0.157 | 0.069 |
| VAL753 | 0.000 | 0.000 | 0.000 | 0.000 | 0.000 | 0.000 | 0.000 | 0.000 | 0.183 | 0.151 | 0.195 | 0.187 | 0.218 | 0.147 | 0.125 |
| MET754 | 0.000 | 0.306 | 0.127 | 0.105 | 0.000 | 0.000 | 0.000 | 0.180 | 0.259 | 0.085 | 0.251 | 0.233 | 0.028 | 0.135 | 0.161 |
| GLY755 | 0.239 | 0.153 | 0.052 | 0.206 | 0.000 | 0.265 | 0.270 | 0.141 | 0.095 | 0.117 | 0.186 | 0.057 | 0.185 | 0.146 | 0.225 |
| ILE756 | 0.039 | 0.248 | 0.035 | 0.226 | 0.000 | 0.000 | 0.204 | 0.000 | 0.146 | 0.279 | 0.266 | 0.091 | 0.051 | 0.130 | 0.176 |
| VAL757 | 0.000 | 0.000 | 0.000 | 0.003 | 0.000 | 0.000 | 0.000 | 0.000 | 0.120 | 0.103 | 0.062 | 0.000 | 0.000 | 0.000 | 0.000 |
| GLY758 | 0.280 | 0.191 | 0.247 | 0.049 | 0.162 | 0.108 | 0.187 | 0.000 | 0.126 | 0.158 | 0.187 | 0.368 | 0.234 | 0.214 | 0.238 |
| GLY759 | 0.040 | 0.176 | 0.189 | 0.199 | 0.074 | 0.112 | 0.097 | 0.049 | 0.124 | 0.070 | 0.224 | 0.153 | 0.072 | 0.258 | 0.352 |
| VAL760 | 0.000 | 0.000 | 0.000 | 0.193 | 0.148 | 0.000 | 0.000 | 0.000 | 0.036 | 0.187 | 0.165 | 0.000 | 0.000 | 0.000 | 0.000 |
| VAL761 | 0.000 | 0.000 | 0.212 | 0.226 | 0.177 | 0.000 | 0.000 | 0.000 | 0.099 | 0.049 | 0.038 | 0.000 | 0.000 | 0.000 | 0.000 |
| SER762 | 0.077 | 0.174 | 0.094 | 0.093 | 0.071 | 0.100 | 0.147 | 0.292 | 0.158 | 0.224 | 0.041 | 0.079 | 0.200 | 0.126 | 0.000 |
| ALA763 | 0.000 | 0.235 | 0.044 | 0.125 | 0.247 | 0.141 | 0.169 | 0.000 | 0.272 | 0.117 | 0.246 | 0.000 | 0.000 | 0.000 | 0.000 |
| VAL764 | 0.000 | 0.000 | 0.000 | 0.278 | 0.000 | 0.000 | 0.000 | 0.000 | 0.000 | 0.153 | 0.000 | 0.000 | 0.000 | 0.000 | 0.000 |
| SER765 | 0.212 | 0.154 | 0.122 | 0.193 | 0.238 | 0.128 | 0.344 | 0.377 | 0.220 | 0.223 | 0.137 | 0.155 | 0.234 | 0.219 | 0.000 |
| GLY766 | 0.214 | 0.209 | 0.209 | 0.216 | 0.250 | 0.168 | 0.274 | 0.187 | 0.148 | 0.142 | 0.334 | 0.140 | 0.095 | 0.015 | 0.000 |
| VAL767 | 0.000 | 0.000 | 0.308 | 0.178 | 0.171 | 0.000 | 0.000 | 0.000 | 0.000 | 0.069 | 0.088 | 0.047 | 0.000 | 0.000 | 0.000 |
| SER768 | 0.000 | 0.000 | 0.000 | 0.000 | 0.150 | 0.000 | 0.000 | 0.000 | 0.036 | 0.080 | 0.151 | 0.136 | 0.353 | 0.000 | 0.000 |
| SER769 | 0.328 | 0.196 | 0.193 | 0.134 | 0.099 | 0.107 | 0.191 | 0.199 | 0.139 | 0.172 | 0.199 | 0.090 | 0.119 | 0.170 | 0.234 |
| PHE770 | 0.170 | 0.000 | 0.168 | 0.000 | 0.000 | 0.305 | 0.000 | 0.000 | 0.000 | 0.340 | 0.150 | 0.152 | 0.148 | 0.000 | 0.163 |
| MET771 | 0.097 | 0.000 | 0.062 | 0.152 | 0.101 | 0.180 | 0.126 | 0.156 | 0.000 | 0.082 | 0.178 | 0.049 | 0.133 | 0.153 | 0.118 |
| SER772 | 0.163 | 0.000 | 0.000 | 0.000 | 0.151 | 0.000 | 0.000 | 0.000 | 0.304 | 0.000 | 0.170 | 0.291 | 0.242 | 0.168 | 0.273 |
| ASN773 | 0.000 | 0.000 | 0.000 | 0.000 | 0.000 | 0.000 | 0.000 | 0.000 | 0.089 | 0.000 | 0.000 | 0.277 | 0.000 | 0.193 | 0.389 |
| PHE775 | 0.062 | 0.000 | 0.000 | 0.155 | 0.065 | 0.113 | 0.000 | 0.000 | 0.000 | 0.000 | 0.194 | 0.000 | 0.000 | 0.000 | 0.246 |
| CC | 0.373 | 0.411 | 0.255 | 0.548 | 0.483 | 0.411 | 0.362 | 0.334 | 0.438 | 0.369 | 0.340 | 0.468 | 0.464 | 0.406 | 0.333 |

**Figure S5**: Reaction Coordinate Coefficients and Stabilities for Postfusion gB. The coefficients between the fusion loop residues (columns) and the membrane proximal region (MPR) residues (rows) are colored on a green scale to denote the magnitude of the coefficients. The bottom row shows the “stability” of each fusion loop residue, which is a measure of how well the reaction coordinate captures a coordinated motion of MPR residues.

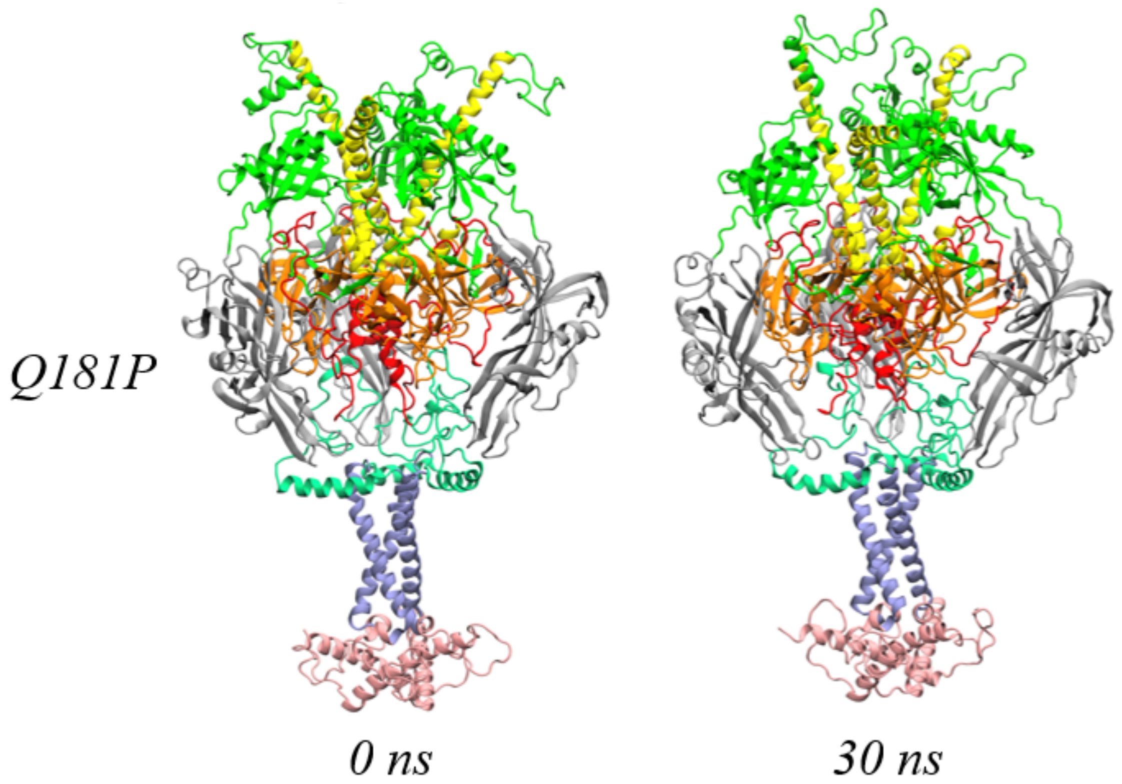


**Figure S6**: Mutant gB Conformations over 30 ns of simulation. A comparison of the starting conformation at 0 ns and the ending conformation at 30 ns for the pre-fusion mutation (Q181P) system reveals a stable binding pose. The different domains are colored as follows: gray for domain I, green for domain II, yellow for domain III, orange for domain IV, red for domain V, light green for membrane-proximal region, purple for transmembrane domain, and pink for cytoplasm tail domain.

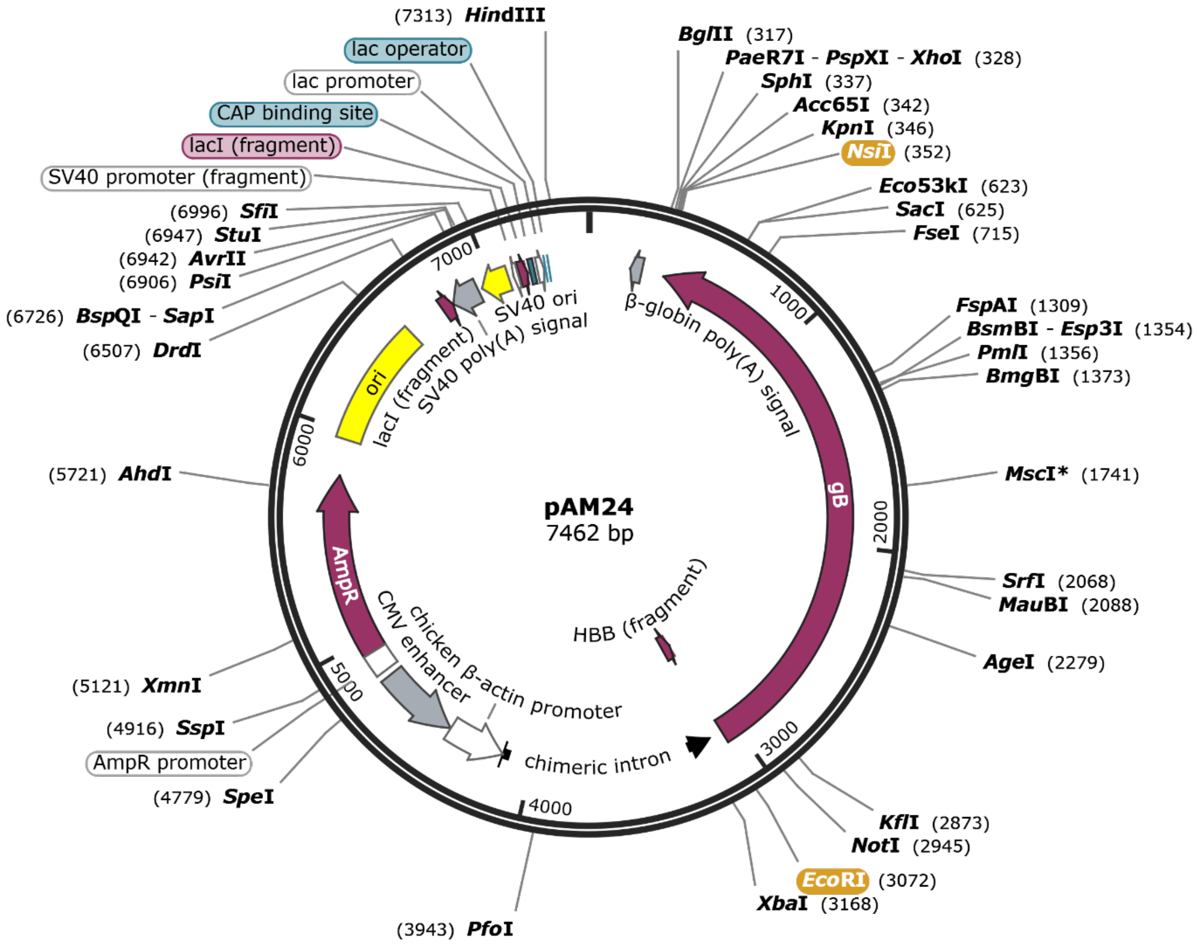


**Figure S7**: Map of plasmid encoding the HSV-1 strain KOS gB gene in a pCAGGS backbone. The plasmid (pAM24) is 7462 base pairs in length and has an AmpR gene for ampicillin resistance, a chicken beta-actin promoter (CβA) with the CMV enhancer that drives the transcription of the gB gene, and multiple cloning sites (MCS) including restriction sites *EcoRI* and *NsiI* (highlighted in orange) upstream and downstream of the gB gene, respectively. An origin of replication (Ori) is located upstream of the gB gene ahead of the SV40 promoter fragment, SV40 polyadenylation signal, and CAP binding site. The plasmid also has a *lacI* gene fragment, lac operator sites, and pBR322 origin of replication. The map highlights the direction of transcription for each feature with arrows indicating the orientation. The plasmid encoding the Q181P gB gene (pAM26) is identical except for the mutation. The image was generated and annotated with SnapGene 7.1.2.

```
HSV-1gB          1 MHQGAPSWGRRWFVVWALLGLTLGVLVASAAPSSPGTPGVAAATQAANGG    50
                   ||||||||||||||||||||||||||||||||||||||||||||||||||
HSV-1gBQ181P     1 MHQGAPSWGRRWFVVWALLGLTLGVLVASAAPSSPGTPGVAAATQAANGG    50

HSV-1gB         51 PATPAPPALGAAPTGDPKPKKNKKPKNPTPPRPAGDNATVAAGHATLREH   100
                   ||||||||||||||||||||||||||||||||||||||||||||||||||
HSV-1gBQ181P    51 PATPAPPALGAAPTGDPKPKKNKKPKNPTPPRPAGDNATVAAGHATLREH   100

HSV-1gB        101 LRDIKAENTDANFYVCPPPTGATVVQFEQPRRCPTRPEGQNYTEGIAVVF   150
                   ||||||||||||||||||||||||||||||||||||||||||||||||||
HSV-1gBQ181P   101 LRDIKAENTDANFYVCPPPTGATVVQFEQPRRCPTRPEGQNYTEGIAVVF   150

HSV-1gB        151 KENIAPYKFKATMYYKDVTVSQVWFGHRYSQFMGIFEDRAPVPFEEVIDK   200
                   |||||||||||||||||||||||||||||.||||||||||||||||||||
HSV-1gBQ181P   151 KENIAPYKFKATMYYKDVTVSQVWFGHRYSPFMGIFEDRAPVPFEEVIDK   200

HSV-1gB        201 INAKGVCRSTAKYVRNNLETTAFHRDDHETDMELKPANAATRTSRGWHTT   250
                   ||||||||||||||||||||||||||||||||||||||||||||||||||
HSV-1gBQ181P   201 INAKGVCRSTAKYVRNNLETTAFHRDDHETDMELKPANAATRTSRGWHTT   250

HSV-1gB        251 DLKYNPSRVEAFHRYGTTVNCIVEEVDARSVYPYDEFVLATGDFVYMSPF   300
                   ||||||||||||||||||||||||||||||||||||||||||||||||||
HSV-1gBQ181P   251 DLKYNPSRVEAFHRYGTTVNCIVEEVDARSVYPYDEFVLATGDFVYMSPF   300

HSV-1gB        301 YGYREGSHTEHTSYAADRFKQVDGFYARDLTTKARATAPTTRNLLTTPKF   350
                   ||||||||||||||||||||||||||||||||||||||||||||||||||
HSV-1gBQ181P   301 YGYREGSHTEHTSYAADRFKQVDGFYARDLTTKARATAPTTRNLLTTPKF   350

HSV-1gB        351 TVAWDWVPKRPSVCTMTKWQEVDEMLRSEYGGSFRFSSDAISTTFTTNLT   400
                   ||||||||||||||||||||||||||||||||||||||||||||||||||
HSV-1gBQ181P   351 TVAWDWVPKRPSVCTMTKWQEVDEMLRSEYGGSFRFSSDAISTTFTTNLT   400

HSV-1gB        401 EYPLSRVDLGDCIGKDARDAMDRIFARRYNATHIKVGQPQYYLANGGFLI   450
                   ||||||||||||||||||||||||||||||||||||||||||||||||||
HSV-1gBQ181P   401 EYPLSRVDLGDCIGKDARDAMDRIFARRYNATHIKVGQPQYYLANGGFLI   450

HSV-1gB        451 AYQPLLSNTLAELYVREHLREQSRKPPNPTPPPPGASANASVERIKTTSS   500
                   ||||||||||||||||||||||||||||||||||||||||||||||||||
HSV-1gBQ181P   451 AYQPLLSNTLAELYVREHLREQSRKPPNPTPPPPGASANASVERIKTTSS   500

HSV-1gB        501 IEFARLQFTYNHIQRHVNDMLGRVAIAWCELQNHELTLWNEARKLNPNAI   550
                   ||||||||||||||||||||||||||||||||||||||||||||||||||
HSV-1gBQ181P   501 IEFARLQFTYNHIQRHVNDMLGRVAIAWCELQNHELTLWNEARKLNPNAI   550

HSV-1gB        551 ASVTVGRRVSARMLGDVMAVSTCVPVAADNVIVQNSMRISSRPGACYSRP   600
                   ||||||||||||||||||||||||||||||||||||||||||||||||||
HSV-1gBQ181P   551 ASVTVGRRVSARMLGDVMAVSTCVPVAADNVIVQNSMRISSRPGACYSRP   600

HSV-1gB        601 LVSFRYEDQGPLVEGQLGENNELRLTRDAIEPCTVGHRRYFTFGGGYVYF   650
                   ||||||||||||||||||||||||||||||||||||||||||||||||||
HSV-1gBQ181P   601 LVSFRYEDQGPLVEGQLGENNELRLTRDAIEPCTVGHRRYFTFGGGYVYF   650
```

```
HSV-1gB          651 EEYAYSHQLSRADITTVSTFIDLNITMLEDHEFVPLEVYTRHEIKDSGLL    700
                     ||||||||||||||||||||||||||||||||||||||||||||||||||
HSV-1gBQ181P     651 EEYAYSHQLSRADITTVSTFIDLNITMLEDHEFVPLEVYTRHEIKDSGLL    700

HSV-1gB          701 DYTEVQRRNQLHDLRFADIDTVIHADANAAMFAGLGAFFEGMGDLGRAVG    750
                     ||||||||||||||||||||||||||||||||||||||||||||||||||
HSV-1gBQ181P     701 DYTEVQRRNQLHDLRFADIDTVIHADANAAMFAGLGAFFEGMGDLGRAVG    750

HSV-1gB          751 KVVMGIVGGVVSAVSGVSSFMSNPFGALAVGLLVLAGLAAAFFAFRYVMR    800
                     ||||||||||||||||||||||||||||||||||||||||||||||||||
HSV-1gBQ181P     751 KVVMGIVGGVVSAVSGVSSFMSNPFGALAVGLLVLAGLAAAFFAFRYVMR    800

HSV-1gB          801 LQSNPMKALYPLTTKELKNPTNPDASGEGEEGGDFDEAKLAEAREMIRYM    850
                     ||||||||||||||||||||||||||||||||||||||||||||||||||
HSV-1gBQ181P     801 LQSNPMKALYPLTTKELKNPTNPDASGEGEEGGDFDEAKLAEAREMIRYM    850

HSV-1gB          851 ALVSAMERTEHKAKKKGTSALLSAKVTDMVMRKRRNTNYTQVPNKDGDAD    900
                     ||||||||||||||||||||||||||||||||||||||||||||||||||
HSV-1gBQ181P     851 ALVSAMERTEHKAKKKGTSALLSAKVTDMVMRKRRNTNYTQVPNKDGDAD    900

HSV-1gB          901 EDDL    904
                     ||||
HSV-1gBQ181P     901 EDDL    904
```

**Figure S8**: Amino acid sequence alignment of HSV-1 strain KOS gB and Q181P gB. Identical residues are indicated by vertical bars (|). Nonidentical residues are indicated by a period (.). The glutamine to proline substitution mutation at residue 181 is bound by a red box. Sequences were analyzed with EMBOSS Stretcher 2.3.0, Pairwise Sequence Alignment.
